# Hierarchical Anchoring–Gating Dictates Specific H4K16 Acetylation by the Human MSL Complex

**DOI:** 10.64898/2026.03.03.709310

**Authors:** Qiang Shi, Yan Zhao, Zhiheng Deng, Jiawei Liang, Shidian Wu, Huasong Ai, Luyang Sun, Lei Liu

## Abstract

Histone acetylation is a versatile post-translational modification essential for diverse biological processes. While nearly all histone acetyltransferases (HATs) act promiscuously on multiple lysine residues, the human male-specific lethal (MSL) acetyltransferase complex mediates strictly site-specific acetylation of histone H4 at lysine 16 (H4K16ac), with its underlying mechanism remaining enigmatic. Here, we leveraged chemical protein synthesis to engineer a semisynthetic nucleosome probe that traps the MSL complex in a catalytically engaged state and determined its 3.12 Å cryo-electron microscopy (cryo-EM) structure, which reveals a hierarchical anchoring–gating mechanism responsible for stringent H4K16 selectivity. The MSL complex adopts a dual-anchoring mode to bind nucleosomal DNA at superhelix location (SHL) 1.5 and the H2A–H2B acidic patch. Three spatially coordinated gating elements (the MSL1 anchor helix, KAT8 gating helix, and KAT8 gating hairpin) then conformationally constrain the H4 N-terminal tail, funneling H4K16 side chain exclusively into the active site while sterically occluding flanking lysines. Targeted mutations disrupting these interfaces impair H4K16ac *in vitro* and in cells, with concomitant defects in chromatin accessibility and transcription. Beyond the monomeric 1:1 complex, cryo-EM analysis identifies a 2:2 MSL–nucleosome assembly, wherein MSL3 dimerization mediates inter-nucleosomal contacts linked to genome-wide H4K16ac patterning. Collectively, our work delineates a residue-to-domain framework in which a chromatin acetylation writer hardwires specific catalysis into nucleosome engagement and scales this precision to higher-order chromatin architecture, providing a mechanistic foothold for understanding aberrant MSL-encoded chromatin programs in diseases.

## Introduction

Histone acetylation is a cornerstone post-translational modification event that laid the conceptual foundation for modern epigenetics^1–3^. By neutralizing the positive charge of lysine side chains on histone tails, acetylation weakens histone–DNA and inter-nucleosomal interactions, reshapes chromatin accessibility, and thereby calibrates transcriptional programs that sustain cellular homeostasis and adaptive responses to stress^4–7^. Extensive studies have shown that nearly all histone acetyltransferases (HATs) exhibit permissive substrate selectivity and act promiscuously on multiple sites, targeting numerous lysines across distinct histone tails to generate combinatorial acetylation landscapes^2,7–11^. Recent studies of nucleosome-bound HATs, including p300^12^ and the NuA4 complex^13,14^, have reinforced the view that acetylation is driven primarily by proximity to the substrate, with HATs either engaging nucleosomal DNA to modify multiple tails without strict positional constraints, or relying on auxiliary subunits to position the catalytic domain for multi-site acetylation on histone H4 (e.g., at K5, K8, K12, and K16). These findings support a prevailing model in which histone acetylation is governed primarily by proximity-driven enzymatic interactions with nucleosomes rather than stringent site-specific recognition. However, against this backdrop of broadly promiscuous acetylation, the acetylation of histone H4 lysine 16 (H4K16ac), deposited by the MSL complex with exquisite positional fidelity on a single lysine site^15–18^, stands out as a conspicuous counterexample. This site-specific acetylation event is evolutionarily conserved and uniquely effective at destabilizing higher-order chromatin folding^19,20^, thereby promoting transcriptional activation, genome maintenance, and cell-cycle regulation^21–33^. Dysregulation of H4K16ac is recurrently observed in multiple cancers (e.g., breast, lung, gastric, hepatocellular carcinomas)^34–44^, and pathogenic mutations in MSL subunits underlie X-linked developmental syndromes^45^.

The MSL complex is a multi-subunit assembly comprising the catalytic MYST-family acetyltransferase KAT8 and the auxiliary subunits MSL1 and MSL3 (**Fig. 1a**)^18,46–48^, which collectively confer nucleosome-dependent acetyltransferase activity and specificity^46,49–51^. Notably, strict H4K16 specificity is not an intrinsic property of the catalytic KAT8 itself. Although KAT8 is also incorporated into other acetyltransferase assemblies (e.g., non-specific lethal complex^31,51^), strict H4K16 selectivity emerges exclusively in the context of the intact MSL complex^51^. Furthermore, while the MSL complex can acetylate free histones promiscuously, robust and selective H4K16 acetylation occurs solely on nucleosomal substrates^51,52^. This dual dependence on both the assembly of the enzymatic complex and the nucleosomal substrate context deviates fundamentally from the prevailing model of proximity-driven, promiscuous acetylation. These biochemical features argue for a distinct, hitherto uncharacterized mechanism that enforces H4K16 acetylation specificity on chromatin, highlighting the need to elucidate how the MSL complex organizes its subunits on the nucleosome to choreograph precise catalytic outcomes.

**Fig. 1.**
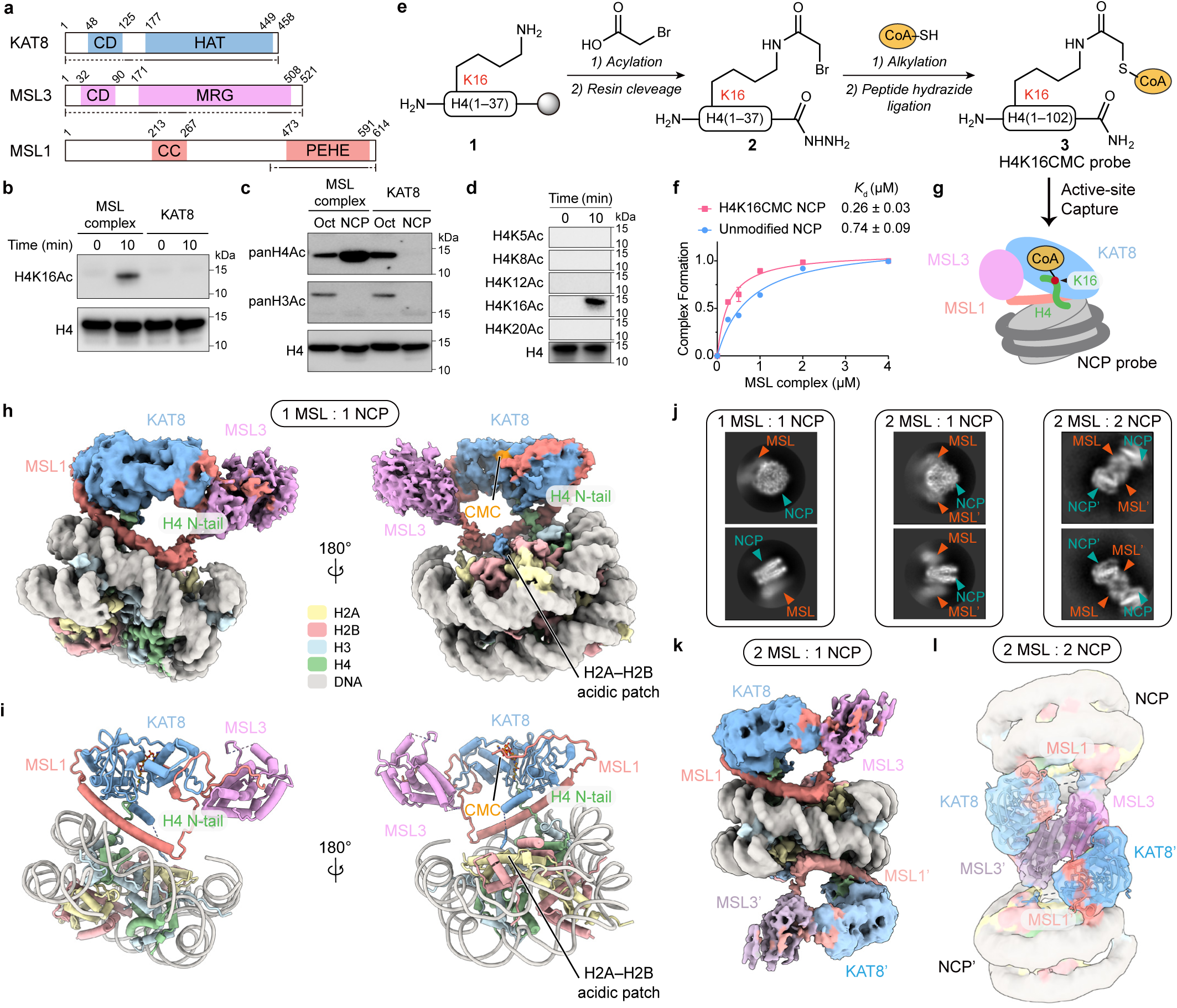
Biochemical and structural basis of H4K16ac by the human MSL complex on the nucleosome. **a**, Domain organization of human KAT8, MSL1, and MSL3. Underlined regions indicate fragments of the reconstituted complexes used in this study, whereas dashed segments denote disordered regions. CD, chromo-barrel domain; HAT, histone acetyltransferase; MRG, *MORF4*-related gene family; CC, coiled coil. **b**, Cr Time-course in vitro acetylation assay comparing H4K16ac by the reconstituted MSL complex and KAT8 alone on nucleosome substrates. **c**, Western blot analysis of bulk histone acetylation on histone octamer and nucleosome substrates by the reconstituted MSL complex or KAT8 alone. **d**, Time-course acetylation assays showing specific modification of H4K16 by the reconstituted human MSL complex on nucleosome substrates. **e**, Semi-synthetic strategy for generating H4K16CMC NCP probe used as a catalytic trapping probe. **f**, Binding affinities of the MSL complex to unmodified and H4K16CMC NCPs, determined by EMSA. *K*_d_ values were obtained by fitting data to One Site-Specific Binding model in GraphPad Prism 9.5.1. Data are represented as mean ± s.d. from n = 3 biological independent experiments. **g**, Schematic of active-site capture of the MSL complex by an H4K16CMC NCP probe. **h**, Cryo-EM density map of the MSL complex bound to H4K16CMC nucleosome in a 1:1 stoichiometry. **i**, Atomic model of the MSL complex bound to H4K16CMC nucleosome in a 1:1 stoichiometry. **j**, Cryo-EM 2D class averages of MSL–NCP complexes at 1:1, 2:1 and 2:2 stoichiometries. NCP electron densities are indicated by green triangles and MSL complex electron densities are indicated by orange triangles. **k**, Cryo-EM density map of the MSL complex bound to H4K16CMC nucleosome in a 2:1 stoichiometry. **l**, Cryo-EM density map of the MSL complex bound to H4K16CMC nucleosome in a 2:2 stoichiometry with the atomic model of fitted MSL complex superimposed as cartoon representations.

To address this knowledge gap, we leveraged chemical protein synthesis to generate a semisynthetic nucleosome probe that enhances the affinity between the MSL complex and nucleosome, stabilizes the catalytically engaged MSL–nucleosome state, and enables structural and mechanistic dissection of nucleosome-dependent H4K16 acetylation targeting. Our cryo-EM structure reveals a hierarchical mechanism: a dual-anchoring mode positions the MSL complex on the nucleosome disk face, while a set of spatially coordinated gating elements—the MSL1 anchor helix, KAT8 gating helix, and KAT8 gating hairpin—geometrically constrain the H4 N-terminal tail. This architecture precisely routes H4K16 into the KAT8 catalytic pocket while excluding neighboring lysine residues. Beyond individual nucleosomes, we observe that cooperative, multivalent engagements among MSL–nucleosome assemblies promote liquid–liquid phase separation, contributing to genome-wide H4K16ac patterning. Collectively, our findings define how the MSL complex encodes the exceptional specificity of H4K16ac and how this precision can be organized across nucleosomes to shape higher-order chromatin behavior.

### Human MSL complex enables nucleosome-dependent specificity for H4K16ac

The human MSL acetyltransferase complex is a multi-subunit assembly that confers both enzymatic activity and substrate specificity for H4K16ac on nucleosomes. The catalytic subunit KAT8 contains an N-terminal chromo-barrel domain (CD) followed by a central histone acetyltransferase (HAT) domain, whereas MSL3 harbors an N-terminal CD domain and a C-terminal *MORF4*-related gene (MRG) domain. MSL1 functions as a scaffold protein, with an N-terminal coiled-coil region and a C-terminal PEHE domain that mediates interactions with both KAT8 and MSL3^18,47,48^ (**Fig. 1a**). We purified a recombinant MSL complex containing full-length KAT8 (1-458), full-length MSL3 (1-521), and a C-terminal fragment of MSL1 (residues 465-614)—a truncation previously shown to retain H4K16 acetylation activity^47,48^ (**Extended Data Fig. 1a**). The purified MSL complex was homogeneous and catalytically active, whereas KAT8 alone showed no detectable activity toward nucleosomal substrates (**Fig. 1b**), demonstrating that productive nucleosome acetylation requires the assembled MSL complex. Comparative acetylation assays using histone octamers versus nucleosomes revealed a pronounced dependence on nucleosomal substrate context (**Fig. 1c**). On octamers, the MSL complex acetylated both H3 and H4, indicative of promiscuous activity. In contrast, on nucleosomes, the acetylation activity was restricted to H4 and was substantially enhanced. KAT8 alone acetylated octamers but was inactive on nucleosomes (**Fig. 1c**). Using antibodies specific for individual H4 acetyl lysine residues, we found that the MSL complex acetylated nucleosomes exclusively at H4K16, with no detectable modification at H4K5, K8, K12, or K20 (**Fig. 1d**). Thus, although both KAT8 and the assembled MSL complex are active on octamers, only the assembled complex acquires robust and selective nucleosomal activity. Consistent with this specificity, genetic perturbation of MSL components (MSL1, KAT8, or MSL3) in cells significantly reduced H4K16ac, while acetylation at other major H4 lysines was largely unaffected (**Extended Data Fig. 1b–d**). These results align with previous studies^49–52^ and establish that the MSL complex confers stringent H4K16 specificity in nucleosomal contexts, both *in vitro* and in cells.

### Chemical capture of a catalytically engaged MSL–nucleosome complex

To capture the catalytically engaged MSL-nucleosome complex, we employed a strategy based on the bisubstrate inhibitor carboxymethyl coenzyme A (CMC)^14,53–55^, which occupies the acetyltransferase active site. It was hypothesized that site-specific covalent attachment of CoA to H4K16 would lock the MSL complex onto the nucleosome in a catalytically engaged state. The H4K16CMC nucleosome probe in which a CMC moiety is conjugated to H4K16 was chemically synthesized (**Fig. 1e**). Briefly, the H4 N-terminal segment (residues 1–37) was first prepared with a bromoacetyl handle installed specifically at K16^14,53–55^, followed by chemoselective conjugation to coenzyme A. The resulting fragment was ligated to H4(39–102) via peptide hydrazide ligation^56^ to generate full-length H4 bearing the K16CMC modification. All synthetic intermediates and the histone H4K16CMC were validated by mass spectrometry (**Supplementary Data. 1**). The histone H4K16CMC was then assembled into nucleosome core particles (NCPs) (**Extended Data Fig. 1e**). Quantitative electrophoretic mobility shift assay (EMSA) demonstrated that the MSL complex binds H4K16CMC NCPs with approximately three-fold higher affinity than unmodified NCP (*K*_d_ = 0.26 ± 0.03 μM vs. 0.74 ± 0.09 μM; **Fig. 1f and Extended Data Fig. 1f**).

We performed single-particle cryo-EM analysis of the MSL complex bound to H4K16CMC NCP probe in an active site (**Fig. 1g**). Data processing yielded a reconstruction at 3.12 Å resolution, revealing the MSL–NCP complex in a 1:1 stoichiometry (**Fig. 1h, i and Extended Data Fig. 2, 3**). The structure reveals that the MSL complex binds to one disk face of the nucleosome in a defined orientation: KAT8, MSL1, and MSL3 form an integrated assembly that engages nucleosomal DNA, the histone core, and the H4 N-terminal tail. MSL1 serves as a central architectural scaffold that bridges the catalytic subunit KAT8 and MSL3. An extended α-helix in MSL1 tracks along the nucleosomal DNA, anchoring the complex and spatially organizing KAT8 and MSL3. A flexible loop (residues I544–P584) emanating from the vicinity of DNA SHL 0 links KAT8 and MSL3, orienting the catalytic center above the H4 tail. Additional density over the H2A–H2B acidic patch corresponds to a loop in KAT8. Strikingly, the trajectory of the H4 N-terminal tail enables insertion of the H4K16 side chain into the KAT8 active site, which accommodates the CMC moiety. This spatial arrangement adopts a constrained nucleosome-bound configuration that enables site-specific acetylation of H4K16 (**Fig. 1h, i and Extended Data Fig. 3**).

Notably, during cryo-EM data processing, we observed multiple MSL–NCP assemblies with distinct stoichiometries: 1:1, 2:1, and 2:2 (**Fig. 1j and Extended Data Fig. 2, 3**). In the 2:1 complex, the additional MSL complex binds to the nucleosome via the same mode as in the 1:1 assembly (**Extended Data Fig. 3k**), preserving the overall binding interface (**Fig. 1k**). In contrast, the 2:2 assembly forms a distinct higher-order structure: two MSL-NCP complexes associate in a pseudo-symmetric, antiparallel arrangement, with each MSL complex bound to its respective nucleosome as observed in the 1:1 state (**Fig. 1l**). In this 2:2 configuration, MSL3 subunits from opposing complexes dimerize via an α-helical interface, while simultaneously contacting DNA on the partner nucleosome—thereby establishing an MSL-mediated inter-nucleosomal architecture (**Fig. 1l**).

### An MSL1 α-helix specifies nucleosomal engagement at DNA SHL 1.5

Cryo-EM analysis reveals that the MSL complex engages nucleosomal DNA via an extended α-helix of MSL1 (residues D500–K541), which contacts the DNA backbone near SHL 1.5 (**Fig. 2a**). Electrostatic surface analysis further demonstrates complementary charge distribution at this interface: the MSL1 helix presents a basic surface that faces the negatively charged DNA phosphate backbone, while residues R516 and R519 within the helix align along the phosphate backbone to form direct contacts that stabilize the complex on the nucleosome (**Fig. 2b, c**). To test the functional contribution of this interface, we generated MSL complexes carrying charge-reversal mutations at the DNA-contacting residues (R516E or R519E) and assessed activity *in vitro*. Both mutants exhibited an approximately 80% reduction in catalytic activity compared to the wild-type complex (**Fig. 2e**). We next evaluated this interface in cells using rescue experiments in *MSL1* knockout (KO) cells. Re-expression of wild-type MSL1 restored global H4K16ac levels, whereas the R516E or R519E mutants failed to do so (**Fig. 2f**). To assess chromatin-level consequences of disrupting the MSL1–DNA anchoring interface, we performed cleavage under targets and tagmentation (CUT&Tag) to profile H4K16ac occupancy and assay for transposase-accessible chromatin with sequencing (ATAC-seq) to measure chromatin accessibility in *MSL1* KO cells reconstituted with wild-type or mutant MSL1. Wild-type MSL1 restored prominent promoter-proximal H4K16ac enrichment and increased chromatin accessibility around transcription start sites (TSSs), whereas the anchoring-helix mutants did not re-establish these features (**Fig. 2j, k**). Consistent with the heatmaps, metagene analyses revealed a strong TSS-centered gain in both H4K16ac and ATAC signal upon wild-type rescue that was markedly blunted in cells expressing R516E or R519E mutants (**Fig. 2j, k**). These results demonstrate that the MSL1–DNA anchoring interface is essential for H4K16ac deposition and the establishment of accessible chromatin states.

**Fig. 2.**
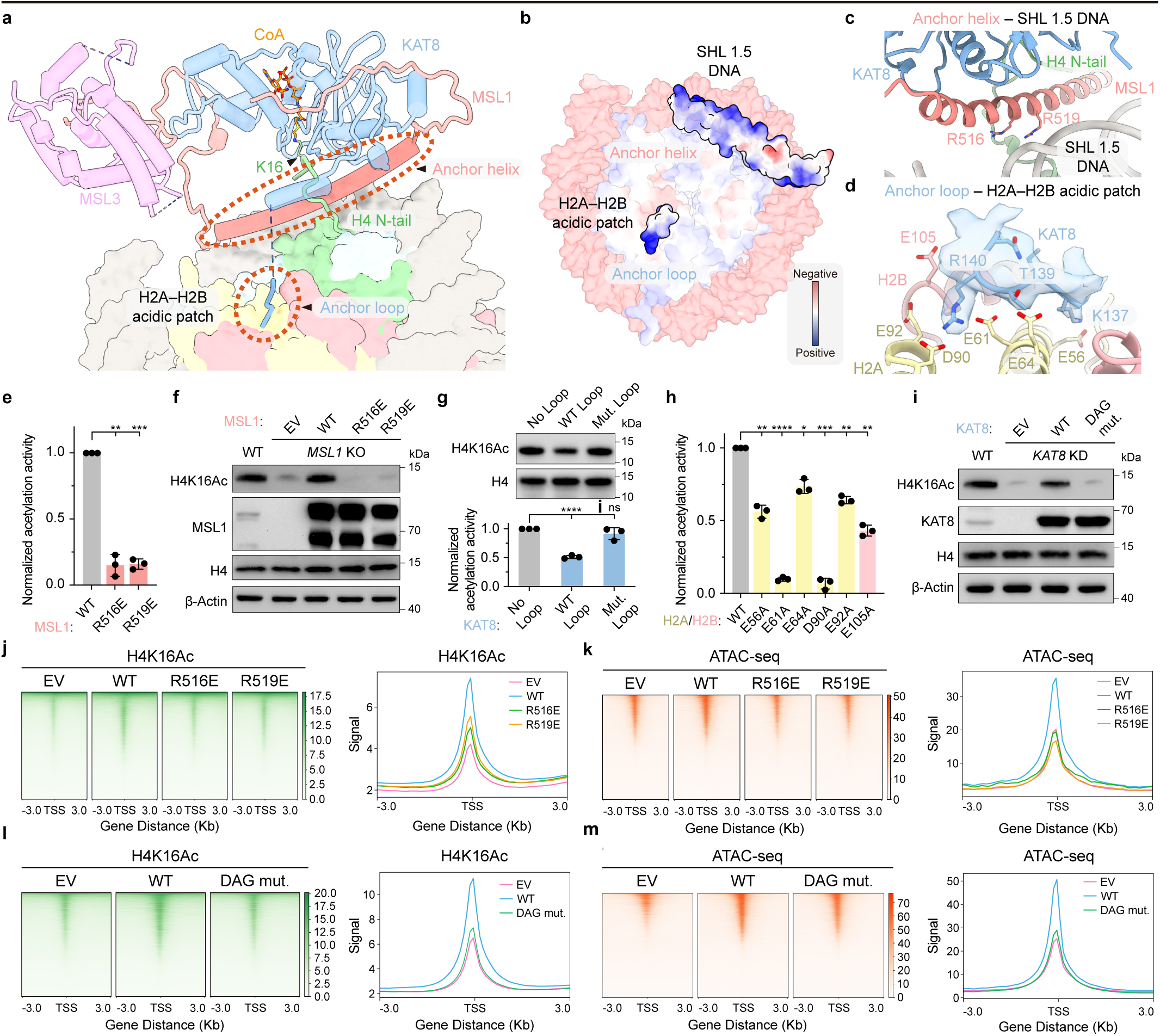
Anchor-mediated nucleosome engagement by the human MSL complex. **a**, Atomic model of the MSL–NCP complex in a 1:1 stoichiometry, represented with the MSL1 anchor helix and KAT8 anchor loop positioned adjacent to SHL 1.5 DNA and the H2A–H2B acidic patch, respectively, and the H4 N-terminal tail oriented toward KAT8 catalytic pocket. **b**, Electrostatic surface representation of the nucleosome, highlighting the SHL 1.5 DNA region and the H2A–H2B acidic patch engaged by the MSL1 anchor helix and KAT8 anchor loop through electrostatic complementarity. **c**, Enlarged view of the MSL1 anchor helix–DNA interface at SHL 1.5, showing positively charged MSL1 residues R516 and R519 contacting the DNA backbone. **d**, Close-up view of local cryo-EM density shows the KAT8 anchor loop positioned near acidic residues on H2A and H2B, defining a charge-complementary interface at the H2A–H2B acidic patch. **e**, Quantification of *in vitro* acetylation activity of MSL complexes containing the indicated MSL1 anchor helix constructs. A two-sample, two-tailed Student’s t-test was employed to calculate P values; **p < 0.01, and ***p < 0.001 Data are represented as mean ± s.d. from n = 3 biological independent experiments. **f**, Western blot analysis of H4K16ac levels in *MSL1* KO cells expressing the indicated MSL1 anchor helix constructs. **g**, Western blot analysis and quantification of *in vitro* acetylation activity of MSL complexes in the absence and presence of KAT8 WT loop or Mut. loop. A two-sample, two-tailed Student’s t-test was employed to calculate P values; ****p < 0.0001 and ns means not significant. Data are represented as mean ± s.d. from n = 3 biologically independent experiments. **h**, Quantification of *in vitro* acetylation activity of MSL complexes on the nucleosomes carrying H2A–H2B acidic patch mutations. A two-sample, two-tailed Student’s t-test was employed to calculate P values; *p < 0.05, **p < 0.01, ***p < 0.001, and ****p < 0.0001. Data are represented as mean ± s.d. from n = 3 biologically independent experiments. **i**, Western blot analysis of H4K16ac levels in *KAT8* KD cells expressing the indicated KAT8 anchor loop constructs. **j**, Heatmaps and average signal profiles of H4K16ac CUT&Tag signal in *MSL1* KO cells expressing the indicated MSL1 anchor helix constructs. **k**, Heatmaps and average signal profiles of ATAC-seq signal in *MSL1* KO cells expressing the indicated MSL1 anchor helix constructs. **l**, Heatmaps and average signal profiles of H4K16ac CUT&Tag signal in *KAT8* KD cells expressing the indicated KAT8 anchor loop constructs. **m**, Heatmaps and average signal profiles of ATAC-seq signal in *KAT8* KD cells expressing the indicated KAT8 anchor loop constructs.

### A KAT8 anchor loop secures nucleosomal positioning via the acidic patch

Cryo-EM analysis uncovers a second anchoring interface between the MSL complex and the nucleosome, mediated by a loop in KAT8 (residues K137–R140) that is positioned above the H2A–H2B acidic patch (**Fig. 2a, d**). This interaction is supported by electrostatic complementarity: the KAT8 loop presents a positively charged patch that aligns with the negatively charged H2A–H2B acidic patch (**Fig. 2b**). At the residue level, K137, T139, and R140 within the KAT8 anchor loop are positioned to contact the acidic patch (**Fig. 2d**).

To validate the functional importance of this interface, we performed *in vitro* biochemical assays. Addition of a synthetic peptide corresponding to the wild-type KAT8 loop peptide (residues S128–N150) effectively competed with the intact MSL complex for H2A–H2B acidic patch binding, reducing nucleosomal acetylation activity by approximately 50%. In contrast, a peptide containing the triple substitutions (K137D, T139A and R140G) showed no inhibitory effect (**Fig. 2g**). Complementarily, mutations within the H2A–H2B acidic patch itself—particularly substitution of core acidic residues E61 and D90—caused an approximately twofold greater impairment of MSL complex–mediated H4K16 acetylation than mutations at peripheral positions (**Fig. 2h**).

We further evaluated the KAT8–acidic patch interface using rescue experiments in *KAT8* knockdown (KD) cells. Re-expression of wild-type KAT8 robustly restored H4K16ac, whereas expression of the DAG mutant (K137D/T139A/R140G) resulted in a markedly reduced rescue activity (**Fig. 2i**). Notably, the R140G substitution in the DAG mutant corresponds to a cancer-associated variant identified in tubular stomach adenocarcinoma^57^. To evaluate chromatin-level effects, we performed H4K16ac CUT&Tag and ATAC-seq in *KAT8* KD cells expressing wild-type KAT8 or the DAG-mutant KAT8. Wild-type KAT8 restored promoter-proximal enrichment of H4K16ac and the associated accessibility landscape, whereas the DAG mutant failed to re-establish these genome-wide profiles (**Fig. 2l, m**). Consistent with these chromatin phenotypes, RNA-seq analysis of MSL complex-regulated genes showed that the DAG mutant only partially restored transcriptional output compared with wild-type KAT8 (**Extended Data Fig. 5**). Together, these results demonstrate that engagement of the H2A–H2B acidic patch by the KAT8 anchor loop is required for productive nucleosome association and chromatin-level establishment of H4K16ac.

### Triple-gating architecture dictates the H4K16ac specificity by the human MSL complex

Building on the dual-anchoring mode—wherein MSL1 engages nucleosomal DNA and KAT8 contacts the H2A–H2B acidic patch—these bipartite interactions stably position the MSL complex to the nucleosome disk face and place the catalytic center of KAT8 directly above the H4 N-terminal tail (**Fig. 2a, 3a**). Close inspection of the MSL–NCP complex reveals three spatially coordinated structural elements in the vicinity of H4K16: the MSL1 gating helix (residues R506–R533), the KAT8 gating helix (residues D160–T174) and the KAT8 gating hairpin (residues K257–E281). Each element contacts the H4 N-terminal tail, and together they constrain its trajectory, effectively inserting the tail into the KAT8 catalytic pocket for acetylation (**Fig. 3a**).

**Fig. 3.**
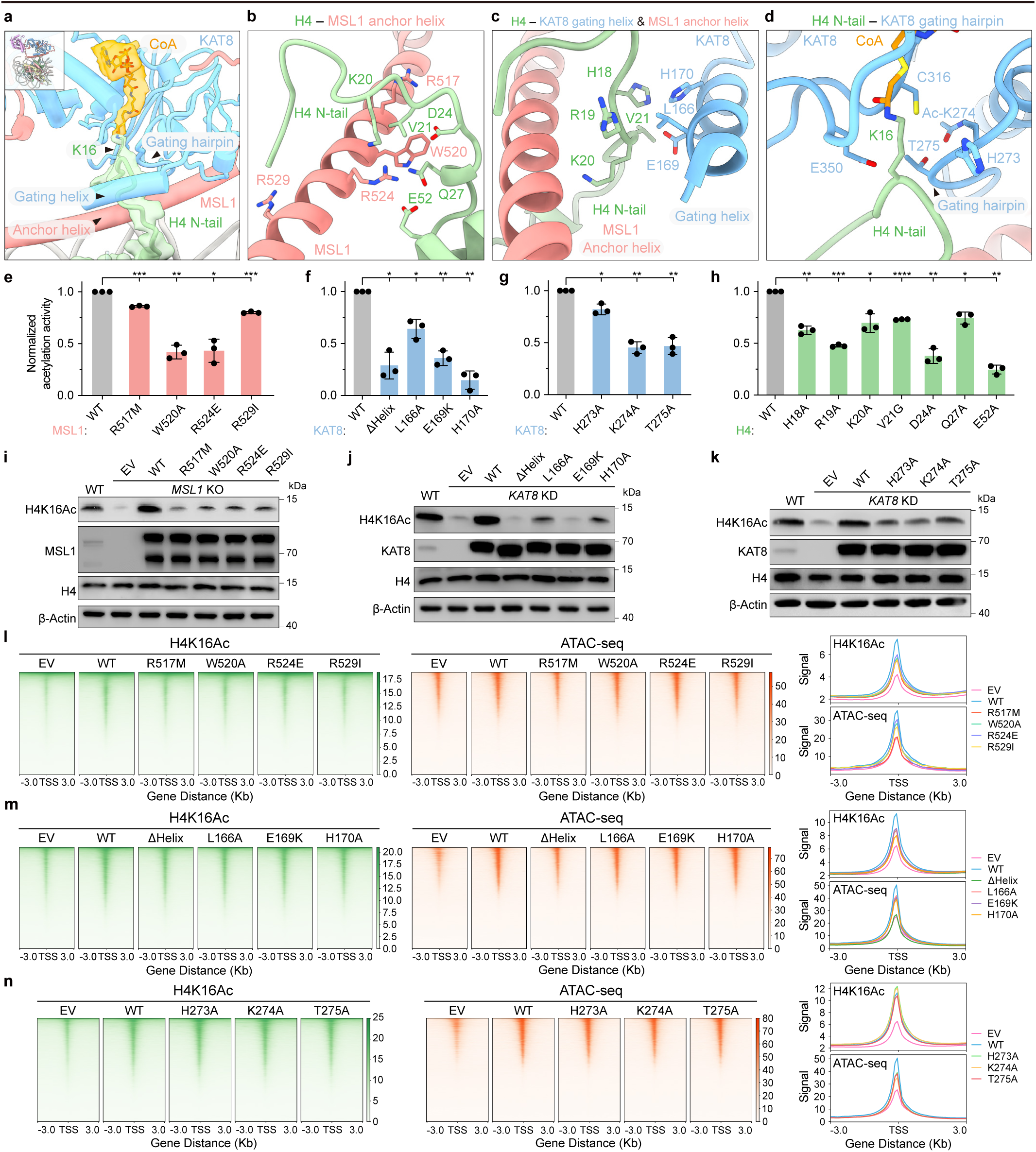
Spatial confinement of the H4 N-terminal tail by MSL1 anchor helix and KAT8 gating helix and gating hairpin underlies precise H4K16ac. **a**, Cryo-EM density map of CoA-conjugated H4 illustrating spatial triple gating mode of the H4 N-terminal tail by MSL1 and KAT8 structural elements. **b**, Close-up view highlighting interactions between the H4 N-terminal tail and the MSL1 anchor helix. **c**, Close-up view showing coordinated contacts between the H4 N-terminal tail, the KAT8 gating helix and the MSL1 anchor helix. **d**, Close-up view of the H4 N-terminal tail positioned adjacent to the KAT8 gating hairpin and catalytic pocket. **e**, Quantification of in vitro acetylation activity of MSL complexes containing the indicated MSL1 anchor helix constructs. **f**, Quantification of in vitro acetylation activity of MSL complexes containing the indicated KAT8 gating helix constructs. **g**, Quantification of in vitro acetylation activity of MSL complexes containing the indicated KAT8 gating hairpin constructs. **h**, Quantification of in vitro acetylation activity of MSL complexes on nucleosomes containing the indicated H4 N-terminal tail variants. A two-sample, two-tailed Student’s t-test was employed in (**g**) to (**j**) to calculate P values; *p < 0.05, **p < 0.01, ***p < 0.001, and ****p < 0.0001. Data are represented as mean ± s.d. from n = 3 biological independent experiments. **i**, Western blot analysis of H4K16ac levels in *MSL1* KO cells expressing the indicated MSL1 anchor helix constructs. **j**, Western blot analysis of H4K16ac levels in *KAT8* KD cells expressing the indicated KAT8 gating helix constructs. **k**, Western blot analysis of H4K16ac levels in *KAT8* KD cells expressing the indicated KAT8 gating hairpin constructs. **l**, Heatmaps and average signal profiles of H4K16ac CUT&Tag signal and ATAC-seq signal in *MSL1* KO cells expressing the indicated MSL1 constructs. **m**, Heatmaps and average signal profiles of H4K16ac CUT&Tag signal and ATAC-seq signal in *KAT8* KD cells expressing the indicated KAT8 gating helix constructs. **n**, Heatmaps and average signal profiles of H4K16ac CUT&Tag signal and ATAC-seq signal in *KAT8* KD cells expressing the indicated KAT8 gating hairpin constructs.

Beyond anchoring the MSL complex to nucleosomal DNA near SHL 1.5, the MSL1 anchoring helix contributes an additional gating function: its H4-facing surface engages the H4 N-terminal tail through conserved residues including R524 (oriented toward H4 E52 and Q27), W520 (adjacent to H4 D24), and additional stabilizing residues R517 and R529, forming complementary contacts that restrict the conformational freedom of the proximal H4 tail segment (**Fig. 3b**).

Opposite the MSL1 anchoring helix, the KAT8 gating helix (residues D160–T174) forms a narrow interaction interface with the H4 N-terminal region, where residues L166, E169, and H170 of the helix are juxtaposed to H4 residues H18 and V21, constraining the rotational flexibility of the H4 tail as it approaches the catalytic center (**Fig. 3c**). Sequence alignment across metazoans reveals high conservation of this KAT8 gating helix (**Extended Data Fig. 4a**), and AlphaFold3-based modeling predicts a similar spatial arrangement among vertebrate KAT8 orthologs; notably, structural superposition reveals that the corresponding helix in Drosophila KAT8 occupies a discernibly shifted position relative to the catalytic core, which may reflect lineage-specific architectural constraints imposed by the fly dosage-compensation complex that incorporates RNA components and auxiliary recognition modules^18,46^ (**Extended Data Fig. 4b**). Comparison across representative human histone acetyltransferases further shows that this gating helix is unique to KAT8 and absent from other human HAT family enzymes associated with promiscuous multi-site acetylation (**Extended Data Fig. 4d**).

Adjacent to these gating features and close to the KAT8 catalytic center, the gating hairpin further sculpts a confined catalytic microenvironment around H4K16, guiding the H4K16 side chain into a configuration aligned with the catalytic cysteine C316 and the general base E350, as corroborated by cryo-EM density of the H4K16–CoA moiety deeply inserted into the active site (**Fig. 3d**). Notably, our cryo-EM reconstruction reveals clear electron density for an acetyl group at K274 (**Extended Data Fig. 4c**), consistent with prior studies^58–60^ establishing K274 (and its orthologs in MYST-family acetyltransferases) as a strictly conserved autoacetylation site essential for activity. Within the gating hairpin, key residues—H273, Ac-K274, and T275—cooperatively stabilize H4K16 engagement and restrict alternative conformations of the H4 tail (**Fig. 3d**).

Together, these three spatially organized gating elements—the MSL1 anchoring helix, KAT8 gating helix, and KAT8 gating hairpin—collectively constitute a triple-gating architecture. This spatial arrangement constrains the H4 N-terminal tail to a defined trajectory towards the KAT8 active site. Structurally, it favors substrate engagement with a single lysine residue while precluding access to alternative sites, enabling the MSL complex to exert precise, nucleosome-dependent control over acetylation specificity via structure-mediated substrate routing, beyond the intrinsic enzymatic activity of KAT8 alone.

### Functional validation of the hierarchical triple-gating architecture in H4K16ac

To test the functional relevance of the triple-gating architecture, we performed *in vitro* nucleosome acetylation assays using MSL complexes harboring mutations in individual gating elements, alongside mutations in H4 tail residues that mediate interactions with these gates. Perturbation of the H4-interacting surface of the MSL1 anchoring helix—including alanine substitution of W520 (W520A), charge reversal at R524 (R524E), and cancer-associated variants R517M and R529I—resulted in reduced nucleosomal acetylation activity relative to the wild-type complex (**Fig. 3e**). Truncation of the KAT8 gating helix (ΔHelix) or substitution of its H4-interfacing residues (L166A, E169K, H170A) substantially diminished acetylation (**Fig. 3f**). Similarly, substitution of key residues in the KAT8 gating hairpin (H273A, Ac-K274A, and cancer-associated variant T275A^57^) also reduced activity (**Fig. 3g**). In parallel, mutations in H4 tail residues that contact the gating elements (H18A, R19A, K20A, V21G, D24A, Q27A, E52A) each led to decreased H4K16ac *in vitro* (**Fig. 3h**), supporting a distributed interaction network that stabilizes the routed H4 tail trajectory.

We next evaluated the triple-gating elements through rescue experiments in *MSL1* KO or *KAT8* KD cells, combined with genome-wide profiling. In *MSL1* KO cells, re-expression of wild-type MSL1 restored global H4K16ac levels, whereas reintroduction of R517M, W520A, R524E, or R529I mutants failed to do so (**Fig. 3i**). Consistent with these effects, H4K16ac CUT&Tag and ATAC-seq revealed that these MSL1 mutants impaired H4K16ac enrichment and chromatin accessibility around TSSs relative to wild-type MSL1 (**Fig. 3l**). In *KAT8* KD cells, wild-type KAT8 re-expression robustly restored H4K16ac, but the gating helix truncation (Δhelix) or interface mutants (L166A, E169K, H170A) did not (**Fig. 3j**). These KAT8 gating helix mutants also exhibited markedly reduced H4K16ac occupancy and accessibility, accompanied by only partial restoration of MSL complex-regulated transcriptional programs (**Fig. 3m and Extended Data Fig. 5**). In contrast, KAT8 gating hairpin mutants (H273A, Ac-K274A, T275A) retained partial capacity to re-establish H4K16ac occupancy and accessibility in *KAT8* KD cells (**Fig. 3n**) but still failed to fully restore H4K16ac levels (**Fig. 3k**) and transcriptional output (**Extended Data Fig. 5**). Together, these results demonstrate a hierarchical contribution of the triple-gating elements to nucleosome-dependent H4K16ac enrichment and its transcriptional consequences.

### MSL3-mediated inter-nucleosomal interactions drive condensate formation and H4K16ac regulation

Beyond the mechanistic insights gained from the 1:1 MSL–nucleosome complex, cryo-EM classification also identified a distinct assembly with 2:2 stoichiometry (**Fig. 1j, l and Extended Data Fig. 2**). In this higher-order configuration, two MSL–nucleosome complexes associate in a pseudo-symmetric, antiparallel arrangement. While each MSL complex maintains the same nucleosome-binding mode as the 1:1 state, inter-complex coupling is specifically mediated by MSL3 (**Fig. 4a**). Specifically, a basic loop (RRKR, residues 193–196) within MSL3 engages the DNA backbone near SHL 6.0 of the opposing nucleosome (**Fig. 4b**), and an α-helical segment (residues T208–F223) forms a stable dimerization interface between MSL3 subunits from adjacent complexes (**Fig. 4c**). These dual MSL3-mediated interactions—with both inter-nucleosomal DNA and opposing MSL3 subunit—establish the structural basis for connecting neighboring nucleosomes.

**Fig. 4.**
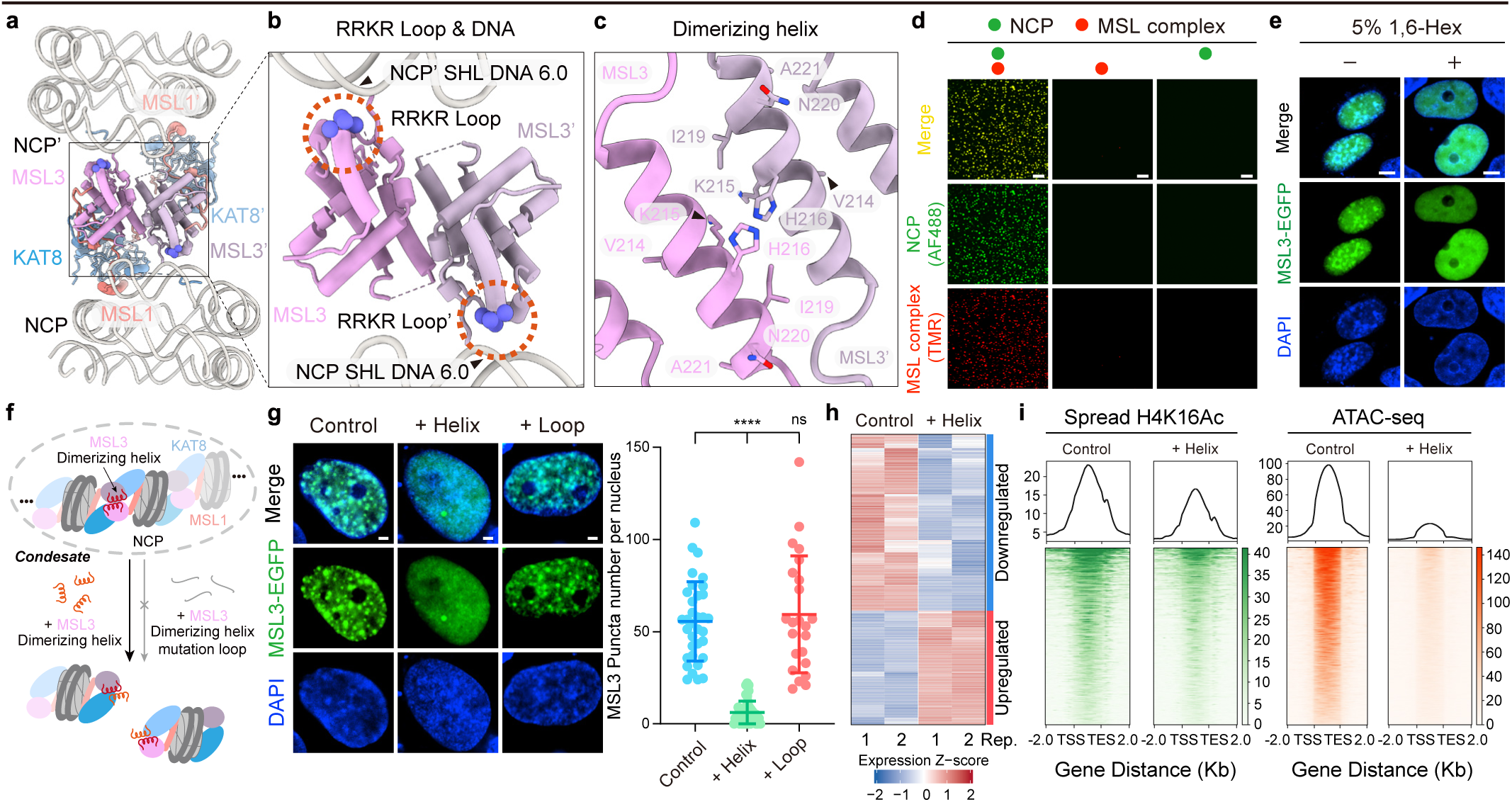
MSL3 dimerization promotes phase separation of MSL–nucleosome assemblies with downstream chromatin consequences. **a**, Cryo-EM structure of MSL–nucleosome assemblies in a 2:2 stoichiometry which two MSL complexes are simultaneously engaged with two nucleosomes. MSL3 is positioned at the interface between adjacent MSL complexes, forming an inter-complex contact that links neighboring nucleosomes. **b**, A basic RRKR (residues 193–196) engages SHL 6.0 DNA of the opposing nucleosome. **c**, Close-up view of the MSL3 dimerizing helix, highlighting residues that define the dimer interface. **d**, *In vitro* condensate formation of MSL–nucleosome assemblies upon the co-presence of NCP and the MSL complex. Scale bar, 20 μm. **e**, Representative fluorescence images showing nuclear MSL3 puncta formed upon overexpression of MSL3–EGFP in cells treated with or without 5% 1,6-Hex. Scale bar, 5 μm. **f**, Schematic illustration of chemical perturbation targeting MSL3 dimerization to modulate phase separation. **g**, Left, representative immunofluorescence images of nuclear MSL3 puncta formed upon overexpression of MSL3–EGFP in cells without treatment or treated with the MSL3 dimerizing-helix peptide or mutant loop peptide. Right, quantification of nuclear MSL3 puncta per nucleus; n = 40 cells without treatment, n = 42 cells with MSL3 dimerizing-helix peptide treatment, and n = 22 cells with mutant loop peptide treatment. A two-sample, two-tailed Student’s t-test was employed to calculate P values; ****p < 0.0001 and ns means not significant. Data are represented as mean ± s.d. Scale bar, 2 μm. **h**, Heat maps of transcriptional changes of control cells and cells treated with MSL3 dimerizing-helix peptide (two replicates each). **i**, Heatmaps and average signal profiles of the spread H4K16ac (left) and corresponding ATAC-seq signal (right) across gene bodies from transcription start sites to transcription end sites in control cells and cells treated with MSL3 dimerizing-helix peptide.

To explore the functional relevance of this inter-nucleosomal coupling, we investigated whether the 2:2 configuration contributes to phase separation, a process increasingly implicated in chromatin regulation^61–67^. In reconstituted *in vitro* systems, purified MSL complexes and nucleosomes cooperatively assembled into liquid-like condensates, whereas neither component alone formed detectable assemblies (**Fig. 4d**). Condensate formation was concentration-dependent and was disrupted by 1,6-hexanediol (1,6-Hex, a reagent that disrupts weak interactions) and elevated ionic strength (**Extended Data Fig. 6a–c**). Live imaging showed dynamic fusion of droplets, and fluorescence recovery after photobleaching (FRAP) assays showed partial recovery of both nucleosome– and MSL-associated fluorescence (**Extended Data Fig. 6d, e**), consistent with the dynamic molecular exchange typical of liquid-like condensates. In cellular contexts, both ectopically expressed MSL3–EGFP and endogenous MSL3 formed discrete nuclear puncta that were sensitive to 1,6-Hex treatment (**Fig. 4e and Extended Data Fig. 6g**); FRAP analysis of MSL3–EGFP validated their dynamic behavior (**Extended Data Fig. 6f**), and endogenous MSL3 puncta co-localized with MSL1 and KAT8 (**Extended Data Fig. 6h**), verifying these are intact, functional MSL complexes. Notably, the MSL3 interfaces mediating inter-complex coupling—including the RRKR basic loop and α-helical dimerization segment—are evolutionarily conserved from Drosophila to mammals (**Extended Data Fig. 7a**). To directly test the contribution of MSL3 dimerization to condensate formation, we designed a short peptide derived from the MSL3 α-helical dimerization region to selectively perturb phase separation (**Fig. 4f**). This wild-type peptide effectively disrupted condensate formation both *in vitro* and in cells, whereas a control peptide—where key helical residues (ILE 209–211, VKH 214–216, INA 219–221) were replaced with glycine (**Extended Data Fig. 7b**)—had no such effect (**Fig. 4g and Extended Data Fig. 7c, d**), supporting that MSL3-mediated dimerization is directly responsible for driving phase separation.

To dissect the chromatin-level consequences of this dimerization, we performed H4K16ac CUT&Tag profiling of H4K16ac and MSL3 occupancy, ATAC-seq, and RNA-seq in control cells and cells treated with the MSL3 dimerizing helix peptide (residues 208–223, TILESYVKHFAINAAF). Compared with controls, peptide-treated cells exhibited coordinated decreases in H4K16ac and MSL3 occupancy, diminished chromatin accessibility, and widespread downregulation of genes regulated by the MSL complex (**Fig. 4h and Extended Data Fig 8a–f**). Genomic annotation revealed that a substantial fraction of H4K16ac (63.0%), MSL3 (62.4%), and ATAC-seq (43.3%) peaks were promoter-proximal (**Extended Data Fig 8b–d**), consistent with a transcription-linked role of the MSL complex. Functional enrichment analysis of differentially expressed genes associated with H4K16ac-enriched loci showed strong overrepresentation of cell-cycle-related pathways, including mitotic progression and chromosome segregation (**Extended Data Fig 8h**), linking condensate disruption to proliferative transcriptional programs. Beyond global attenuation, condensate disruption preferentially weakened extended domains of H4K16ac enrichment. Genome-wide analysis showed marked reduction of both spread H4K16ac signals and chromatin accessibility across broad genomic regions (**Fig. 4i**), and genome browser views illustrated concomitant weakening of H4K16ac, MSL3 binding, and ATAC-seq signals across extended chromatin domains (**Extended Data Fig 8g**). Together, these results support a model in which MSL3-mediated inter-nucleosomal coupling promotes condensate formation that helps sustain H4K16ac patterning and chromatin accessibility at both promoter-proximal sites and broader acetylation domains.

## Discussion

Most histone acetyltransferases possess broad substrate tolerance and can acetylate multiple lysine residues across histone tails, a feature that generally precludes strict, site-selective modification^2,7–11^. In contrast, the human MSL acetyltransferase complex is specialized in depositing the highly stringent and consequential H4K16ac mark^18,49–51^. Our work reveals that this specificity arises from a geometric constraint imposed by the MSL–nucleosome assembly. Central to this geometry is a hierarchical anchoring-gating architecture: dual anchoring via MSL1 (to DNA at SHL 1.5) and KAT8 (to the H2A–H2B acidic patch) fixes the complex on the nucleosome disk and positions the KAT8 catalytic center above the H4 N-terminal tail. Built upon this anchored configuration, three coordinated gating elements—the MSL1 anchor helix, KAT8 gating helix, and KAT8 gating hairpin—then collectively constrain the H4 tail’s trajectory and conformational flexibility, channeling only K16 into the active site. This multi-component architecture converts acetylation from a proximity-driven reaction to a topology-dependent process, explaining the strict nucleosomal context requirement for H4K16ac (**Fig. 5**). Consequently, specific H4K16ac is restricted to the nucleosomal context, providing a mechanistic basis for the MSL complex’s unusually stringent site selectivity in cells.

**Fig. 5.**
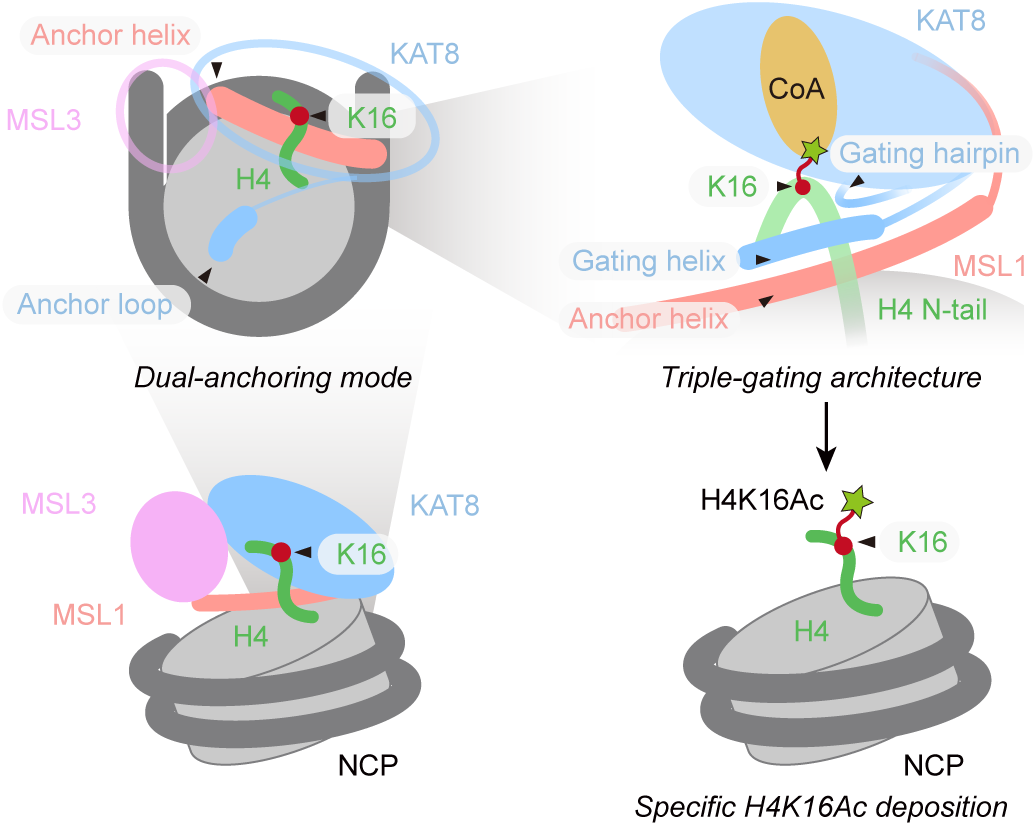
A hierarchical anchoring–gating mechanism for site-specific H4K16 acetylation by the MSL complex. Schematic model illustrating a dual-anchoring mode and a triple-gating architecture by the MSL complex that spatially confine the H4 N-terminal tail, precisely positioning H4K16 toward the KAT8 catalytic pocket to enable site-specific H4K16 acetylation on individual nucleosomes.

Our findings highlight distinct structural logics employed by distinct acetyltransferases. Unlike the DNA-centered, largely non-specific binding mode of p300^12^—wherein the single-subunit enzyme associates with nucleosomes primarily through DNA interactions via its bromodomain and HAT domain (**Extended Data Fig 9a, d**), enabling promiscuous acetylation of all four core histones. And the position-based targeting of NuA4^13,14^, where non-catalytic Epl1 subunit anchors the catalytic Esa1 to the nucleosome disk face via DNA and the H2A–H2B acidic patch without enforcing fixed a fixed substrate trajectory (consequently resulting in multi-lysine acetylation within the H4 tail) (**Extended Data Fig 9b, e**), the human MSL complex adopts a uniquely integrated strategy that couples nucleosome recognition to precise catalysis. Specifically, the MSL complex leverages coordinated contributions from both non-catalytic (MSL1-mediated DNA binding) and catalytic (KAT8 anchor loop engagement with the H2A–H2B acidic patch) subunits to establish a fixed spatial registry with the H4 N-terminal tail, while successive gating interactions progressively narrow the H4 tail’s conformational landscape, thereby permitting productive acetylation only at K16 (**Extended Data Fig 9c, f**). This integrated design enables the MSL complex to achieve exquisite lysine-level specificity for H4K16, distinct from the broad or multi-site modification profiles of p300 and NuA4. These comparisons extend our mechanistic understanding of nucleosome-targeted acetylation, highlighting how architectural constraints within enzyme–nucleosome assemblies can dictate substrate specificity.

Consistent with this model, disrupting the anchoring or gating interactions of MSL has profound chromatin-level consequences. Disease-associated variants linked to the MSL pathway map to the very interfaces that secure nucleosome engagement and substrate routing. For instance, cancer-associated variants include MSL1 variants on the H4-tail–interacting surface (R517M, identified in pancreatic adenocarcinoma and R529I, identified in colon adenocarcinoma), a KAT8 anchor-loop substitution at the H2A–H2B acidic patch (R140G, identified in tubular stomach adenocarcinoma), and a KAT8 gating-hairpin variant adjacent to the catalytic pocket (T275A, identified in uterine endometrioid carcinoma)^57^. Such perturbations would be expected to compromise nucleosome positioning and/or destabilize the constrained H4-tail trajectory required for selective H4K16ac (**Fig. 2, 3**). In this light, loss of H4K16ac is a recurrent feature of cancers and developmental disorders harboring MSL mutations^34–45^, a likely direct consequence of breaking the structural constraints needed for MSL’s function. The diminished H4K16ac and attendant chromatin accessibility defects observed in these disease contexts can be understood as a failure to maintain the nucleosomal anchoring–gating architecture, linking localized interface disruptions to broad epigenomic and transcriptional dysregulation.

At the chromatin scale, our findings suggest how precise nucleosome-level targeting by MSL can be amplified into broader chromatin domain effects. The 2:2 MSL–nucleosome assemblies we identified provide a structural basis for local reinforcement of acetylation (**Fig. 4**). Such inter-nucleosomal coupling and condensate-like organization could act in concert with other features of active chromatin to promote regional acetylation. For example, H3K36 methylation (a mark associated with transcription elongation) has been linked to recruitment of the MSL complex to gene bodies^68–71^. These and other permissive chromatin cues could bias the local chromatin environment toward productive H4K16 acetylation. Rather than operating as a processive spreading mechanism, the collective multivalent interactions in the MSL clusters are likely to increase the dwell time and local concentration of the complex at specific genomic loci, thereby favoring the establishment of H4K16ac on multiple neighboring nucleosomes.

Technically, our study highlights how chemical biology approaches can render fleeting enzyme–substrate encounters structurally tractable. By leveraging a semisynthetic nucleosome probe in which a bisubstrate inhibitor^14,53–55^, carboxymethyl coenzyme A (CMC), is site-specifically conjugated to H4K16, we stabilized the MSL–nucleosome complex in a catalytically engaged state and trapped a transient intermediate of the acetylation reaction. This chemical capture strategy enabled direct visualization of the H4 N-terminal tail during active MSL engagement. Structural superposition of H4 densities revealed that, in the catalytically trapped complex, continuous density extends beyond K16 toward residues preceding D24 (H4 density shown in green), indicating stabilization of an ordered and productive tail conformation. In contrast, on the opposite side, lacking MSL complex engagement, density for the H4 N-terminal region upstream of D24 is markedly weaker and fragmented (H4 density shown in purple), consistent with increased conformational flexibility (**Extended Data Fig 10**). These observations indicate that bisubstrate-mediated catalytic trapping stabilizes a productive H4 tail trajectory, highlighting the power of chemical biology to resolve transient enzyme–substrate interactions.

## Acknowledgments

We thank the National Natural Science Foundation of China (T2488301, to L. L.), National Key R&D Program of China (2022YFC3401500, to L. L.), The Ministry of Science and Technology of China (2021YFA1300603, to L. S.), National Natural Science Foundation of China (22137005, 22227810, to L. L.; 32501108, to H. A.; U24A200281, 32350020, 32370620, to L. S.), New Cornerstone Investigator Program (to L. L.), Peking University Medicine plus X Pilot Program-Key Technologies R&D Project (2024YXXLHGG006, L. S.), Shanghai Natural Science Foundation (25ZR1402193, H. A.), Shanghai Frontiers Science Center of Drug Target Identification and Delivery (ZXWH2170101, H. A.) and Shanghai Jiao Tong University 2030 Initiative (WH510363004/013, H. A.). We acknowledge the Tsinghua University Branch of China National Center for Protein Sciences (Beijing) for cryo-EM data screening and collection. We thank Yanli Zhang and Bingyu Liu (Imaging Core Facility of the Protein Research Center for Technology Development, Tsinghua University) for technical assistance for fluorescence imaging using Nikon AXR NSPARC confocal microscope and Imaris software. We thank Jingjing Wang at Cell Biology Facility, Center of Biomedical Analysis, Tsinghua University for the confocal imaging and analysis.

## Author contributions

Conceptualization: Q.S., H.A., L.S. and L.L. Design of experiments: Q.S., Y.Z., Z.D., H.A., L.S. and L.L. Probe synthesis: Q.S. Cloning and protein purification: Q.S., Z.D., J.L. and S.W. Cryo-EM sample preparation: Q.S. and Z.D. Cryo-EM data processing and model building: H.A., Q.S. and Z.D. Biochemical assays: Q.S. LLPS–related assays: Q.S. and Y.Z. Cellular assays: Y.Z. and Q.S. Genomics sample preparation and data analysis: Y.Z. Data interpretation: S.Q. and Y.Z. Visualization: S.Q., Y.Z. and Z.D. Funding acquisition: L.L., L.S., and H.A. Supervision: L.L., L.S., and H.A. Writing – original draft: Q.S. and Y.Z. Writing – review & editing: L.L., L.S., H.A., Q.S., Y.Z. and Z.D.

## Data and materials availability

The MSL–nucleosome cryo-EM model and map reported in this paper have been deposited in the Protein Data Bank (PDB) with accession numbers 23FT (the 1:1 cryo-EM structure of MSL-H4K16CMC nucleosome) and Electron Microscopy Data Bank (EMDB) EMD-68933 (the 1:1 cryo-EM structure of MSL-H4K16CMC nucleosome), EMD-68934 (the 2:1 cryo-EM structure of MSL-H4K16CMC nucleosome) and EMD-68936 (the 2:2 cryo-EM structure of MSL-H4K16CMC nucleosome). The raw sequencing data for CUT&Tag, ATAC-seq and RNA-seq analyses presented in this paper have been deposited in Gene Expression Omnibus with the accession number GSE318456, GSE318125 and GSE318124, respectively. Newly created materials from this study may be requested from the corresponding authors.

## Methods

### Plasmid construction

Human MSL1 (Uniprot entry Q68DK7, truncation, amino acid residues 465–614), KAT8 (Uniprot entry Q9H7Z6, full length, amino acid residues 1–458), and MSL3 (Uniprot entry Q8N5Y2, full length, amino acid residues 1–521) were synthesized with sequence optimization for *Escherichia coli* recombinant expression by GenScript Biotech (Nanjing, China). The cDNA of human MSL1, KAT8 and MSL3 were synthesized with sequence optimization for Hela cell overexpression by GenScript Biotech (Nanjing, China).

For *E. coli* recombinant expression, MSL1 was cloned into a pGEX-6P-1 vector with an N-terminal GST–HRV3C-tag. KAT8 and MSL3 were cloned into a pET28b(+) vector with an N-terminal His–SUMO-tag. Individual human core histone cDNAs (H2A, H2B, H3 C96S/C110S, and H4) were cloned into a pET22b(+) vector.

For Hela cell overexpression, MSL1, KAT8 and MSL3 were cloned into a pcDNA3.1 expression plasmid with an N-terminal FLAG-tag. For fluorescence imaging, the MSL3 coding sequence was cloned upstream of the EGFP sequence in the FUGW-EGFP vector, generating an N-terminal MSL3–EGFP fusion construct.

Point mutations or truncations were generated by standard site-directed PCR mutagenesis or by homologous recombination.

### Protein expression and purification

The human MSL complex was recombinantly expressed in *E. coli* and purified by sequential affinity chromatography followed by size-exclusion chromatography.

For KAT8 expression, plasmids encoding His–SUMO–tagged KAT8 were transformed into *E. coli* BL21(DE3) cells (Transgene). Cultures were grown in Luria–Bertani (LB) medium containing 50 μg mL⁻¹ kanamycin at 37 °C to an optical density at 600 nm (OD_600_) of ∼0.8, followed by induction with 0.5 mM isopropyl β-D-thiogalactopyranoside (IPTG) and incubation overnight at 18 °C. Cells were harvested by centrifugation and resuspended in ice-cold KAT8 lysis buffer (20 mM HEPES, 150 mM NaCl, 1 mM DTT, pH 7.5) supplemented with 1 mM phenylmethylsulfonyl fluoride (PMSF). Cell disruption was achieved by sonication, and lysates were clarified by centrifugation at 12,000 rpm for 40 min at 4 °C. The supernatant was applied to a Ni–NTA affinity column, and His–SUMO–KAT8 was eluted with buffer containing 300 mM imidazole. Eluted fractions were dialyzed overnight at 4 °C against KAT8 dialysis buffer (20 mM HEPES, 150 mM NaCl, 1 mM DTT), during which the His–SUMO tag was removed by Ulp1 proteases.

In parallel, plasmids encoding GST–HRV3C–MSL1 and His–SUMO–MSL3 were co-transformed into *E. coli* BL21(DE3) cells (Transgene). Cells were cultured in LB medium supplemented with 50 μg mL⁻¹ ampicillin and kanamycin at 37 °C to an OD_600_ of ∼0.8, and protein expression was induced with 0.4 mM IPTG followed by overnight incubation at 18 °C. Harvested cells were lysed by sonication in ice-cold MSL lysis buffer (20 mM HEPES, 400 mM NaCl, 1 mM DTT, pH 7.5) supplemented with 1 mM PMSF. Clarified lysates were loaded onto a Ni–NTA column, and the GST–HRV3C–MSL1/His–SUMO–MSL3 complex was eluted using buffer containing 300 mM imidazole. Eluted fractions were dialyzed overnight at 4 °C against MSL dialysis buffer (20 mM HEPES, 400 mM NaCl, 1 mM DTT), during which the His–SUMO tag on MSL3 was cleaved by Ulp1 proteases. On the following day, the MSL1/MSL3 complex was combined with purified KAT8 and incubated with Glutathione Beads 4FF (Lableads) for 2 h at 4 °C. After extensive washing, assembled MSL complexes were eluted using GST elution buffer containing 30 mM reduced glutathione. The eluate was subsequently treated overnight with HRV3C proteases to remove the GST tag.

Finally, the MSL complex was further purified by size-exclusion chromatography (SEC) using a Superdex 200 10/300 GL column (GE Healthcare) equilibrated in SEC buffer (20 mM HEPES, 150 mM NaCl, pH 7.5) on an ÄKTA system. Peak fractions corresponding to the intact MSL complex were pooled, concentrated, and stored at −80 °C.

Individual human core histones (H2A, H2B, H3 C96S/C110S, and H4) were recombinantly expressed in *E. coli* BL21(DE3) cells (Transgene) and purified as previously described^72^.

### Peptide synthesis

Solid-phase peptide synthesis (SPSS): Peptides were synthesized by Fmoc-based SPSS using a CEM LibertyBlue™ automated microwave peptide synthesizer. Amino acid couplings were performed using *N,N′*-diisopropylcarbodiimide (DIC) and ethyl cyano(hydroxyimino)acetate (Oxyma) as activating reagents. For each coupling cycle, standard amino acids were coupled under microwave irradiation at 95 °C for 3 min, whereas cysteine and histidine residues were coupled at 50 °C for 3 min to minimize side reactions. Fmoc deprotection was carried out after each coupling step using piperidine-based solutions (10% (v/v) piperidine in DMF). Peptides were assembled on Rink amide resin or hydrazide-functionalized resin, depending on the peptide construct. Upon completion of peptide chain assembly, peptides were cleaved from the resin using a trifluoroacetic acid (TFA)–based cleavage cocktail consisting of thioanisole, water, 1,2-ethanedithiol, triisopropylsilane, and TFA in a ratio of 2:2:1:2:33 (v/v). Cleavage reactions were performed at room temperature for 2–3 h with gentle agitation. Cleavage solutions were concentrated under a nitrogen stream, and crude peptides were precipitated with cold diethyl ether, collected by centrifugation, and dissolved in dilute aqueous TFA. Final purification was performed by reverse-phase high-performance liquid chromatography (RP-HPLC), peptide identity was confirmed by mass spectrometry, and purified peptides were lyophilized.

Installation of the auxiliary handle^53–55^: To enable site-specific installation of an auxiliary handle at H4K16, peptides H4(1-37) were synthesized by Fmoc-based SPSS. During peptide synthesis, the α-amino group of the first residue was protected with a tert-butoxycarbonyl protecting group (Boc), while the ε-amino group of Lys16 was protected with an allyloxycarbonyl protecting group (Alloc), allowing for orthogonal deprotection and selective modification at H4K16. After completion of peptide synthesis, selective removal of the Alloc protecting group from the H4K16 side chain was carried out on resin. Briefly, Pd(PPh_3_)_4_ (60 mg) was dissolved in dichloromethane (DMC) (2 mL), and phenylsilane (PhSiH_3_, 600 μL) was prepared separately. The peptide resin was washed sequentially with DMF and DCM (three times each) and swollen in DCM prior to addition of Pd(PPh_3_)_4_ and PhSiH_3_. The reaction mixture was agitated at 30 °C for 12 h. After deprotection, the resin was washed extensively with DCM and DMF, followed by treatment with a DMF solution containing sodium diethyldithiocarbamate (DDTC; 200 mg in 40 mL DMF), which was added three times, with agitation at 30 °C for 15 min each, to remove residual palladium. The resin was subsequently washed with DMF prior to further modification. Following Alloc deprotection, installation of the auxiliary handle was achieved by on-resin acylation of the H4K16 ε-amine with excess bromoacetic acid (Tokyo Chemical Industry (TCI)) using DIC and Oxyma as coupling reagents in DMF. The reaction was allowed to proceed with gentle agitation at 37 °C overnight. After completion, the resin was thoroughly washed with DMF to remove excess reagents. The modified peptide was subsequently cleaved from the solid resin using a TFA–based cleavage cocktail, followed by purification by RP-HPLC and verification by mass spectrometry.

Covalent attachment of CoA via alkylation: Covalent conjugation of coenzyme A (CoA) was achieved via alkylation of the H4K16-bromoacetyl peptide. H4(1–37)-NHNH_2_-K16-bromoacetyl peptide was dissolved in 6 M guanidine hydrochloride to a final peptide concentration of 5 mM, and the pH was adjusted to 8.0. CoA (Bidepharm) was added at a threefold molar excess (final concentration, 15 mM) relative to the peptide, and the pH was readjusted to 8.0 as needed. The reaction mixture was incubated at 37 °C overnight in the dark. Reaction mixture was purified by RP-HPLC, followed by verification by mass spectrometry.

Peptide hydrazide ligation^56^: Peptide ligation via a hydrazide intermediate was carried out under acidic activation conditions followed by thiol-mediated coupling. The N-terminal H4(1–37)-NHNH_2_-K16CMC fragment was prepared at a final concentration of 1 mM in a ligation buffer composed of 6 M guanidine hydrochloride, adjusted to pH 2.3. The solution was precooled in an ice–salt bath (−20 °C), after which sodium nitrite was introduced at a tenfold molar excess. The reaction was maintained at −20 °C for 30 min with continuous stirring to effect conversion of the hydrazide intermediate. Subsequently, 4-mercaptophenylacetic acid (MPAA) was added at a fortyfold molar excess, and the mixture was briefly incubated on ice before adjustment of the pH to 5.0. The C-terminal H4(38-102) fragment was then introduced, followed by further adjustment of the reaction pH to 6.5. Ligation was allowed to proceed at 37 °C overnight. The ligation mixture was subjected to purification by RP-HPLC. Product identity and integrity were confirmed by mass spectrometry, and purified ligation products were collected and lyophilized.

### Nucleosome reconstitution

Recombinant human histone octamers were assembled from purified core histones^73^. Individual histones H2A, H2B, H3, and H4, or their corresponding variants, were combined in equimolar proportions and solubilized in denaturing buffer containing 10 mM HEPES (pH 7.5), 6 M guanidine hydrochloride, 1 mM DTT, and 1 mM EDTA. Octamer formation was achieved by gradual removal of the denaturant through dialysis against high-salt refolding buffer composed of 10 mM HEPES, 2 M NaCl, 1 mM DTT, and 1 mM EDTA (pH 7.5). Assembled histone octamers were isolated by SEC using a Superdex 200 Increase 10/300 GL column equilibrated in refolding buffer.

The Widom 601 nucleosome positioning sequence (147 bp) was generated by restriction enzyme digestion of plasmid templates following established procedures^74^.

The sequence of the 147-bp Widom 601 DNA used in this study is shown below: CTGGAGAATCCCGGTGCCGAGGCCGCTCAATTGGTCGTAGACAGCTCTAGCACCGCTTAAACGCACGTACGCGCTGTCCCCCGCGTTTTAACCGCCAAGGGGATTACTCCCTAGTCTCCAGGCACGTGTCAGATATATACATCCTGT.

Nucleosomes were reconstituted by controlled salt dilution. Purified histone octamers were combined with 147-bp Widom 601 DNA at equimolar ratios in high-salt conditions (2 M NaCl). Assembly was initiated by progressively lowering the ionic strength via dialysis against HE buffer (10 mM HEPES, 1 mM EDTA, pH 7.5) using a peristaltic pump until the final salt concentration reached approximately 0.1 M. Fully assembled nucleosomes were subsequently purified by anion-exchange chromatography using a DEAE column on an ÄKTA system.

### Cryo-EM sample preparation

For cryo-EM analysis, 2.4 μM human MSL complex was combined with 2 μM reconstituted H4K16CMC NCP probe at the indicated molar ratio in SEC buffer (20 mM HEPES, 25 mM NaCl, pH 7.5). The mixture was incubated at 25 °C for 20 min to allow complex formation. To stabilize the complex prior to grid preparation, chemical crosslinking was performed by addition of an equal volume of crosslinking buffer containing 0.3% (v/v) glutaraldehyde prepared in SEC buffer, yielding a final glutaraldehyde concentration of 0.15% (v/v). The reaction was incubated at 25 °C for 10 min and quenched by addition of 2 M Tris-HCl (pH 7.5) to a final concentration of 100 mM. The crosslinked complex was concentrated and further purified by SEC using a Superose 6 Increase 10/300 GL column (Cytiva) equilibrated in SEC buffer. Peak fractions corresponding to the intact MSL–nucleosome complex were pooled and concentrated to approximately 200 ng/μL for cryo-EM grid preparation. For grid preparation, a 3.5 µL of purified complex was applied to glow-discharged Quantifoil Au R1.2/1.3 300-mesh grids (Quantifoil Micro Tools). Grids were incubated at 4 °C under 100% humidity for 60 s, and plunge-frozen into liquid ethane using a Vitrobot Mark IV syntem (FEI/Thermo Fisher Scientific). Prepared grids were stored in liquid nitrogen until data collection.

### Cryo-EM data collection and image processing

Cryo-EM data were collected on a 300 kV Titan Krios transmission electron microscope equipped with a K3 direct electron detector (Gatan) in 1.0979 Å pixel size. Total dose is 50e^−^/Å^2^. 32 frames were collected in each movie, with 2.56s exposure time. Defocus was set to the range of –1.0 to –2.0 μm. The AutoEMation v.2.0^75^ software was employed for automatic data collection. A total of 9,923 micrographs were recorded and subjected to motion correction using MotionCor (version 2-1.2.6)^76^ and contrast transfer function (CTF) estimation using Gctf (version 1.88)^77^. Automated particle picking followed by particle extraction at bin4 yielded 10,365,196 particles. Extracted particles were subjected to two rounds of two-dimensional (2D) classification, resulting in 8,436,568 particles retained for three-dimensional (3D) analysis. An initial round of 3D classification (T = 4) was performed, producing 3,090,746 particles that were further classified to resolve distinct MSL–nucleosome assemblies. Particles corresponding to the MSL–NCP 1:1 assembly were enriched through successive rounds of 3D classification (T = 4). From this analysis, 245,245 particles were retained and refined to generate a 3.12 Å reconstruction of the complete MSL–NCP 1:1 complex. To improve density quality at the MSL–nucleosome interface, focused 3D classification using a soft mask targeting the MOF–MSL1 anchor-helix region was performed without alignment. A focused refinement on the MOF–MSL1 anchor-helix region using the same particle set yielded a locally enhanced map at 3.80 Å resolution. Then they were combined as the composite map in UCSF Chimera X (version 1.10.1)^78^.

Besides, a subset of 174,261 particles was selected and re-extracted at full pixel size for 3D refinement with C2 symmetry, yielding a reconstruction at 3.27 Å resolution after postprocessing corresponding to the MSL–NCP 2:1 assembly.

In parallel, particles corresponding to higher-order assemblies were processed separately. For MSL–NCP 2:2 assembly, iterative 3D classification and refinement identified 12,631 particles, producing a reconstruction at 7.84 Å resolution.

CTF refinement and Bayesian polishing were applied during later stages of processing where indicated. All final resolutions were determined using the gold-standard Fourier shell correlation (FSC = 0.143)^79^ criterion. Image processing was carried out using RELION^80^, and structural visualization was performed using UCSF ChimeraX (version 1.10.1).

### Model building, refinement and validation

Atomic model building of the MSL–nucleosome complex was performed using the final cryo-EM map at 3.12 Å resolution together with locally refined maps where indicated. Initial rigid-body fitting was carried out in UCSF ChimeraX (version 1.10.1) using a combination of predicted and experimentally determined structure including an AlphaFold3-predicted model of MSL–nucleosome complex, the NuA4 HAT–nucleosome structure (PDB 7VVU), a canonical nucleosome structure (PDB 7XD1), the crystal structure of the MSL1–MSL3 subcomplex (PDB 2Y0N), and the MSL1–KAT8 complex (PDB 2Y0M). All components were independently docked into the cryo-EM density. Following initial docking, the individual components were merged into a single molecular model and manually adjusted in Coot. Regions lacking clear density support were trimmed, and side chains with ambiguous density were removed in WinCoot (0.9.8.95)^81^. The N-terminal tail of histone H4 was built manually residue by residue in WinCoot (0.9.8.95), guided by continuous cryo-EM density and stereochemical restraints. Iterative rounds of real-space refinement were carried out in PHENIX (1.20.1)^82^ using secondary-structure and geometry restraints where appropriate. Between refinement cycles, the model was visually inspected and manually corrected to improve local fit. Statistics for model refinement and validation are summarized in **Supplementary Table 1**.

### In vitro acetylation assays

In vitro nucleosomal acetylation assays were performed using reconstituted nucleosomes and purified MSL complex. For standard reactions, MSL complex was mixed with nucleosomes at final concentrations of 20 nM and 0.3 µM, respectively. Acetyl-coenzyme A (Ac-CoA) (Solarbio) was added to a final concentration of 30 µM to initiate the reaction. The total reaction volume was adjusted with assay buffer (20 mM HEPES, 150 mM NaCl, pH 7.5). Reactions were incubated at 30 °C for the indicated times.

For fluorescence-based detection of acetylation, 7-diethylamino-3-(4′-maleimidylphenyl)-4-methylcoumarin (CPM) (Solarbio) was used as a thiol-reactive probe^83–86^. CPM was freshly prepared and added to the reaction at a 1:1 volume ratio relative to the reaction volume to a final concentration of 3 µM after 15 min of incubation. Following CPM addition, reactions were protected from light. Samples were then transferred to a 384-well black solid assay plate, and fluorescence was measured at an excitation wavelength of 392 nm and an emission wavelength of 482 nm using a plate-reading luminometer (BioTek). Negative-control reactions lacking nucleosomes were performed in parallel to correct for baseline signal and MSL autoacetylation. Fluorescence signals were background-subtracted and normalized to the wild-type MSL complex, and data were processed and plotted using GraphPad Prism 9.5.1.

For time-course acetylation assays, reactions were performed for the indicated time and quenched by the addition of 4× SDS loading buffer. Quenched samples were heated at 95 °C for 5 min, separated on 4–12% SDS–PAGE gels (Genscript), and analyzed by western blotting.

For KAT8 loop competition assay, WT or mutant loop peptides (500 µM) were added to the reaction mixtures. Reactions were performed for 10 min and quenched by the addition of 4× SDS loading buffer. Quenched samples were heated at 95 °C for 5 min, resolved on 4–12% SDS–PAGE gels (GenScript), and analyzed by western blotting.

### EMSA

Electrophoretic mobility shift assays were performed to examine binding of the human MSL complex to nucleosomes. Unmodified NCP or H4K16CMC NCP (25 nM) was incubated with the MSL complex prepared as twofold serial dilutions (0–8 µM) in EMSA buffer (10 mM HEPES, 1 mM EDTA, pH 7.5,) at 4 °C for 10 min. After incubation, samples were supplemented with 6× native loading buffer containing 0.25% (w/v) bromophenol blue and 40% (v/v) glycerol, and resolved on a 4.5% native polyacrylamide gel. Electrophoresis was carried out at 120 V in 0.5× TBE buffer (40 mM Tris-HCl, 45 mM boric acid, and 1 mM EDTA, pH 8.0) at 4 °C for 1 h. Gels were stained with SYBR Gold stain (Invitrogen, Thermo Fisher Scientific).

### Generation of fluorescently labeled MSL complex

Fluorescent labeling of the MSL complex was achieved by covalent conjugation of a tetramethylrhodamine (TMR) fluorophore to the catalytic subunit KAT8, followed by reconstitution and purification of the intact complex. Purified KAT8 protein was incubated with Tetramethylrhodamine-5-Maleimide (Invitrogen, Thermo Fisher Scientific) at a molar ratio of fluorophore to protein (1.1:1) at room temperature for 2 h with gentle mixing and protected from light. Following the labeling reaction, excess free fluorophore was removed by desalting to eliminate unreacted tetramethylrhodamine. The resulting TMR-labeled KAT8 was subsequently used for assembly of the MSL complex. Reconstitution and purification of the TMR-labeled MSL complex were performed following the same protocol as used for the unlabeled MSL complex, including affinity purification and SEC. All steps were conducted under low-light conditions to minimize photobleaching. The purified TMR-labeled MSL complex was concentrated, flash-frozen in liquid nitrogen, and stored at −80 °C until further use.

### Phase separation assays in vitro

Phase separation assays in vitro were performed by incubating purified TMR-labeled MSL complexes and Alexa Fluor 488-labeled nucleosomes in phase separation buffer (10 mM HEPES, 1 mM EDTA, pH 7.5) under the indicated experimental conditions, as detailed in the figure legends. Reaction mixtures were gently mixed and incubated at room temperature for 10 min to allow droplet formation. Following incubation, samples were transferred to 384-well glass-bottom plates (Cellvis, P384LB-1.5H) and immediately imaged using a Nikon AXR NSPARC confocal microscope equipped with a 100× oil-immersion objective. All images were acquired using identical acquisition settings within each experiment to ensure comparability across conditions. Droplet formation was quantified using ImageJ.

### Live-cell imaging and quantification of nuclear condensates

For live-cell imaging, HeLa cells transiently transfected with FUGW–EGFP plasmids encoding MSL3 wild type (MSL3-WT) were seeded onto glass-bottom culture dishes (Nest, 801001). Prior to imaging, nuclei were counterstained with Hoechst 33342 (Invitrogen, Thermo Fisher Scientific) (10 µg/mL) for 5 min at room temperature in the dark. Fluorescence images were acquired using a Nikon AXR NSPARC confocal microscope equipped with either a 60× or 100× oil-immersion objective.

Quantification of nuclear condensates was performed using NIS-Elements software. Nuclei were first segmented based on Hoechst staining, and fluorescence intensity thresholds were defined to identify MSL3–EGFP puncta. Condensates were detected and classified according to fluorescence intensity and area, and the number of puncta per nucleus was recorded using the built-in analysis functions. To account for cell-to-cell variability, three independent biological replicates were performed, with multiple cells analyzed per replicate. Data are presented as scatter plots showing the mean ± s.d. from independent experiments.

### FRAP analysis

In vitro FRAP experiments were conducted on phase-separated MSL (6 µM) –NCP (3 µM) assemblies and imaged on a Nikon AXR NSPARC confocal microscope. Circular regions of interest within individual condensates were photobleached using 10% laser power. Fluorescence recovery was recorded at 2 s intervals for 15 min. Pre-bleach images were acquired to establish baseline intensity. Fluorescence signals were background-subtracted and normalized to the pre-bleach values to generate recovery curves.

For live-cell FRAP analysis, HeLa cells transiently expressing MSL3–EGFP were imaged 48 h after transfection using a Nikon AXR NSPARC confocal microscope. Nuclear puncta were selected for photobleaching using 6% laser power. Time-lapse imaging was performed at 2 s intervals for a total duration of 3 min following bleaching. Pre-bleach images were collected for normalization. Fluorescence intensities were background-corrected and normalized to the average pre-bleach signal to quantify recovery kinetics. Multiple puncta from independent cells were analyzed.

### Cell immunofluorescence

Hela cells were washed twice with PBS buffer and fixed with 4% paraformaldehyde (Solarbio) for 20 min at room temperature. After fixation, cells were permeabilized with PBS containing 0.5% Triton X-100 (Yeasen) for 30 min and blocked with QuickBlock^TM^ Blocking Buffer for Immunol Staining (Beyotime) for 1 h. Cells were then incubated with primary antibodies diluted in QuickBlock^TM^ Primary Antibody Dilution Buffer for Immunol Staining (Beyotime) overnight at 4°C, followed by washing five times with PBS containing 0.5% Triton X-100 (5 min each). Cells were subsequently incubated with fluorophore-conjugated secondary antibodies diluted in QuickBlock^TM^ secondary Antibody Dilution Buffer for Immunol Staining (Beyotime) for 1 h at room temperature in the dark. Nuclei were stained with DAPI (Invitrogen, Thermo Fisher Scientific). After final washes five times with PBS containing 0.5% Triton X-100 (5 min each), cells were mounted using Mounting Medium, antifadin (Solarbio) and imaged as indicated.

The antibodies were used as follows: anti-MSL1 (Novus Biologicals, NBP1-90506, 1:200 dilution), anti-MSL3 (Novus Biologicals, H00010943-B01P, 1:100 dilution), anti-KAT8 (Proteintech, 13842-1-AP, 1:200 dilution), YSFluor^TM^ 594 Goat Anti-Mouse IgG (H+L) (Yeasen, 33212ES60, 1:300 dilution), YSFluor^TM^ 488 Goat Anti-Rabbit IgG (H+L) (Yeasen, 33106ES60, 1:300 dilution).

### Peptide treatment

The dimerizing helix and mutant loop peptides were used in both in vitro and celluar assays. Both peptides were applied at a final concentration of 24 µM. For in vitro phase separation assays, peptides were added simultaneously with purified MSL complexes and nucleosomes and incubated for 10 min at room temperature prior to imaging. For live-cell imaging and omics-based analyses, peptides were added directly to the culture medium. Cells were incubated with peptides for 12 h before downstream analyses, including fluorescence imaging and omics-related experimental procedures.

### Establishment of KO and KD cell lines and rescue experiments

Stable *MSL1* KO cell lines were generated using CRISPR–Cas9–mediated genome editing. Stable *KAT8* KD and *MSL3* KD cell lines were established using short hairpin RNA (shRNA). The targeting sgRNA and shRNAs are list in **Supplementary Table 1**.

For lentiviral production, recombinant plasmids encoding sgRNA or shRNA constructs were co-transfected with packaging plasmids (psPAX2 and pMD2.G) into HEK293T cells. Viral supernatants were harvested 48 h after transfection, clarified by filtration, and concentrated by ultracentrifugation. Target cells were transduced with the concentrated viral particles and selected with 1.5 µg mL⁻¹ puromycin beginning 72 h post-transduction. Knockout or knockdown efficiency was verified by western blot analysis.

For rescue experiments, FLAG-tagged MSL1, KAT8, or MSL3 constructs described above were transiently introduced into the corresponding KO or KD cell lines using Lipofectamine 3000 (Invitrogen, Thermo Fisher Scientific) according to the manufacturer’s instructions.

### Western Blot Analysis

For analysis of MSL1, KAT8, and MSL3 expression, total protein lysates were prepared from HeLa cells. Cells were lysed in RIPA buffer supplemented with a protease inhibitor cocktail, and protein concentrations were determined using a BCA assay. Equal amounts of protein were denatured, resolved by SDS–PAGE, and transferred onto nitrocellulose (NC) membranes (Cytiva). Membranes were blocked with 5% (w/v) non-fat milk in TBST (Tris-buffered saline containing 0.1% Tween-20) for 1 h at room temperature and incubated with the indicated primary antibodies overnight at 4 °C. After three washes with TBST, membranes were incubated with horseradish peroxidase (HRP)–conjugated secondary antibodies. Immunoreactive signals were detected using enhanced chemiluminescence (ECL) and visualized on autoradiography film (Tanon).

For histone modification analysis, histones were acid-extracted from HeLa cells as previously described^87^, with minor modifications. Briefly, cells were harvested and resuspended in nuclei extraction buffer (10 mM Tris-HCl, 1 mM KCl, 1.5 mM MgCl_2_, and 1 mM DTT, pH 8.0) supplemented with protease inhibitors. Isolated nuclei were resuspended in 0.2 M H_2_SO_4_, and histones were extracted overnight at 4 °C. Following centrifugation at 16,000 × g for 10 min at 4 °C, the supernatant was collected, neutralized with 0.4 M NaOH, and prepared for western blot analysis.

The antibodies were used as follows: anti-MSL1 (Novus Biologicals, NBP1-90506, 1:2500 dilution), anti-MSL3 (Thermo Fisher Scientific, PA5-28282, 1:1000 dilution), anti-KAT8 (Abcam, ab200660, 1:2000 dilution), anti-H4K16ac (Abcam, ab109463, 1:3000 dilution), anti-H4K5ac (ABclonal, A23080, 1:10000 dilution), anti-H4K8ac (ABclonal, A7258, 1:7000 dilution), anti-H4K12ac (ABclonal, A22754, 1:2000 dilution), anti-H4K20ac (Abcam, ab177191, 1:2000 dilution), anti-panH3Ac (Abcam, ab300641, 1:1000 dilution), anti-panH4ac (Abcam, ab177790, 1:10000 dilution), anti-Histone H4 (PTMab, PTM-1009, 1:2000 dilution), anti-β-actin (Biodragon, B0099, 1:3000 dilution), HRP-Goat Anti-Rabbit (Biodragon, BF03008, 1:5000 dilution), HRP-Goat Anti-Mouse (Biodragon, BF03001, 1:5000 dilution).

### ATAC-seq

ATAC-seq was performed using the Chromatin Profile Kit (Novoprotein, N248) according to the manufacturer’s instructions. Briefly, nuclei were isolated and subjected to tagmentation by incubation with Tn5 transposase at 37 °C for 30 min. Following tagmentation, sequencing adapters were added, and libraries were amplified by PCR. Amplified libraries were purified using DNA clean beads and assessed for concentration and fragment size distribution using Qubit and Bioanalyzer. Libraries were sequenced on an Illumina NovaSeq X Plus platform to generate 150-bp paired-end reads. All experiments were performed with at least two independent biological replicates.

Raw sequencing reads were trimmed using Fastp (v0.23.2; –-detect_adapter_for_pe) and aligned to the human reference genome (hg38) using Bowtie2 (v2.2.5; –-very-sensitive –X 2000). PCR duplicates were removed with Picard MarkDuplicates (REMOVE_DUPLICATES=true). To correct for Tn5 transposase insertion bias, aligned reads were offset by +4 bp for the positive strand and −5 bp for the negative strand. Peaks were identified using MACS2.

Differential chromatin accessibility analysis was performed using DiffBind (v3.3) in R with the edgeR method, applying a false discovery rate (FDR) threshold of <0.05 and an absolute log_2_ fold change >0.58. Peak annotation was carried out using ChIPseeker, with promoter regions defined as ±3 kb relative to transcription start sites (TSSs). De novo motif enrichment analysis was conducted using HOMER (findMotifsGenome.pl; –len 8,10,12). Signal profiles and heatmaps were generated using deepTools.

### CUT&Tag

CUT&Tag experiments were performed using a commercial kit (Novoprotein, N259-YH01) according to the manufacturer’s instructions. Briefly, Concanavalin A–coated magnetic beads (ConA beads) were activated and incubated with washed cells for 15 min at room temperature to immobilize cells. Cell–bead complexes were incubated with the indicated primary antibodies in antibody buffer for 2 h at room temperature with gentle rotation, followed by incubation with secondary antibodies for 1 h. After washing, samples were incubated with 0.04 mM hyperactive pA–Tn5 transposase–adapter complex in Dig-300 buffer for 1 h. Tagmentation was carried out in 300 µL tagmentation buffer at 37 °C for 1 h and terminated by addition of stop buffer. DNA was purified using DNA extraction beads, and sequencing libraries were constructed by adapter ligation followed by PCR amplification. Libraries were quantified and assessed for quality before sequencing on an Illumina NovaSeq X Plus platform to generate 150-bp paired-end reads. All experiments were performed with at least two independent biological replicates.

Paired-end sequencing reads were trimmed using Fastp (v0.23.2) and aligned to the human reference genome (hg38) using Bowtie2 (v2.2.5; –-very-sensitive –X 1000). PCR duplicates were removed using Picard. Peaks were identified using MACS2 (-g hs –B –q 0.05 –f BAMPE). Normalized signal tracks (bigWig files) were generated using deepTools (bamCoverage –-normalizeUsing RPGC) for downstream visualization and analysis.

### RNA-seq

Total RNA was extracted from cultured cells using TRIzol reagent (TransGen Biotech) according to the manufacturer’s instructions. RNA-seq library preparation and sequencing were performed by Berry Genomics. Libraries were generated for the following samples: *MSL1* KO HeLa cells and rescue cell lines (*MSL1* KO + empty vector; *MSL1* KO + MSL1 wild type MSL1; *MSL1* KO + MSL1 R516E; *MSL1* KO + MSL1 R519E; *MSL1* KO + MSL1 R517M; *MSL1* KO + MSL1 W520A; *MSL1* KO + MSL1 R524E; *MSL1* KO + MSL1 R529I); *KAT8* KD HeLa cells and rescue cell lines (*KAT8* KD + empty vector; *KAT8* KD + wild type KAT8; *KAT8* KD + KAT8 DAG; *KAT8* KD + KAT8 ΔHelix; *KAT8* KD + KAT8 L166A; *KAT8* KD + KAT8 E169K; *KAT8* KD + KAT8 H170A; *KAT8* KD + KAT8 H273A; *KAT8* KD + KAT8 K274A; *KAT8* KD + KAT8 T275A). All experiments were performed with at least two independent biological replicates.

Raw sequencing reads were processed using Fastp (--detect_adapter_for_pe) and aligned to the human reference genome (hg38) using HISAT2 (v2.2.1). Gene-level read counts were obtained with featureCounts (v2.0.6; –t exon –g gene_id –p). Differential gene expression analysis was performed using DESeq2 (v1.44.0), with differentially expressed genes defined by an adjusted P value < 0.05 and an absolute log₂ fold change > 0.58. Normalized signal tracks (bigWig files) for visualization were generated using deepTools (bamCoverage –-normalizeUsing BPM).

---

**Extended Data Fig. 1.**
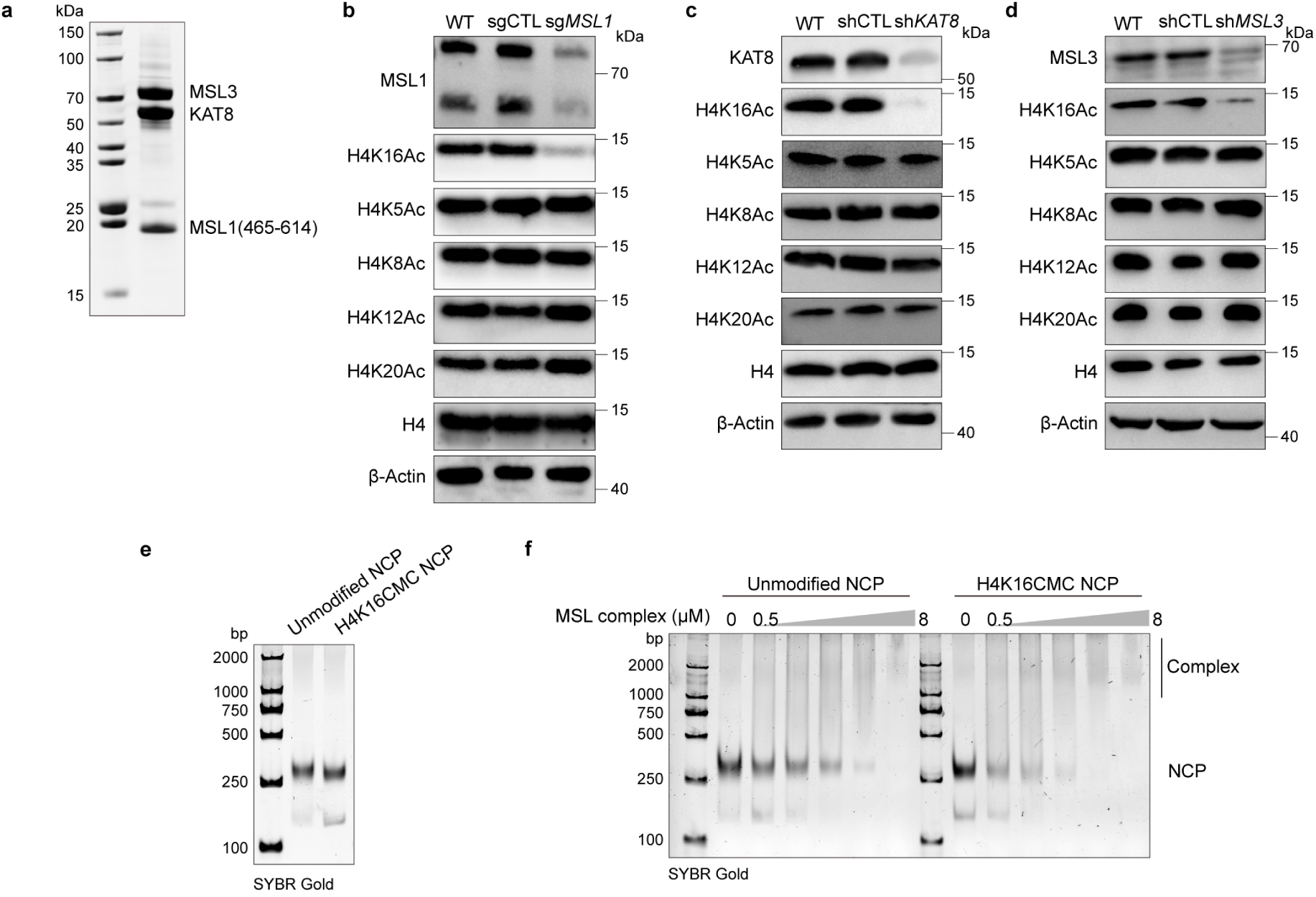
Biochemical and cellular validation of nucleosome-specific H4K16 acetylation by the human MSL complex. **a**, SDS–PAGE analysis of the recombinant human MSL complex used in this study (MSL1(465–614), MSL3 and KAT8). **b**–**d**, Western blot analysis profiling acetylation levels at distinct H4 lysine sites in wild-type cells and cells transduced with control (sgCTL or shCTL), *MSL1*-targeting (sg*MSL1*) sgRNAs, *KAT8*-targeting (sh*KAT8*) shRNAs or *MSL3*-targeting (sh*MSL3*) shRNAs using site-specific H4 acetylation antibodies. **e**, Native PAGE analysis of unmodified and H4K16CMC NCPs stained with SYBR Gold. **f**, EMSA analysis of MSL–nucleosome complex formation with unmodified or H4K16CMC NCPs across increasing concentrations of the MSL complex, revealing enhanced binding of MSL complex to H4K16CMC NCPs.

**Extended Data Fig. 2.**
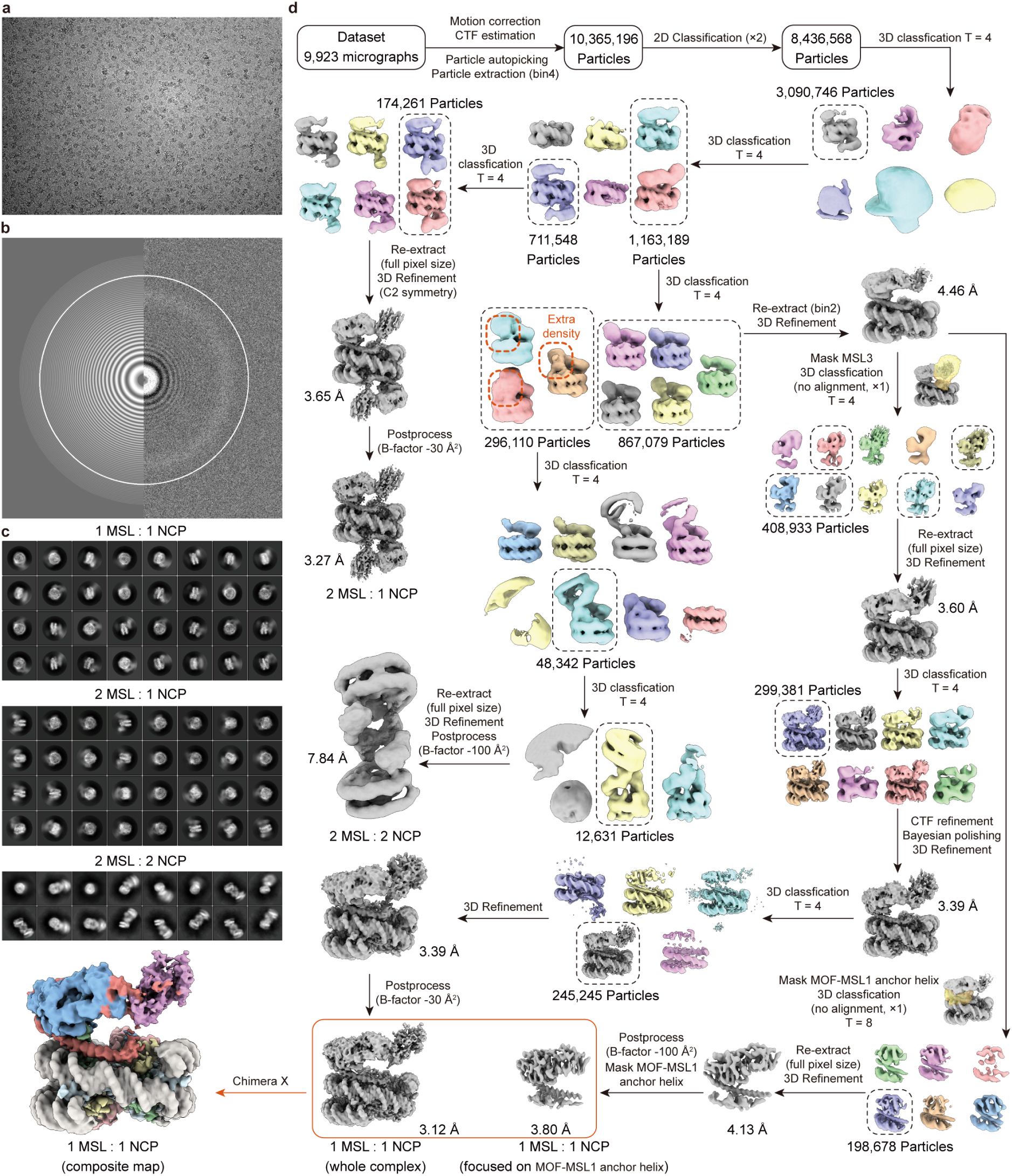
Cryo-EM data processing for MSL–H4K16CMC NCP complex. **a**, Representative micrograph from the dataset. **b**, CTF estimation of the micrograph shown in **a**. **c**, Representative 2D averages for assemblies of MSL**–**NCP in different stoichiometric ratios. **d**, Cryo-EM processing flow chart.

**Extended Data Fig. 3.**
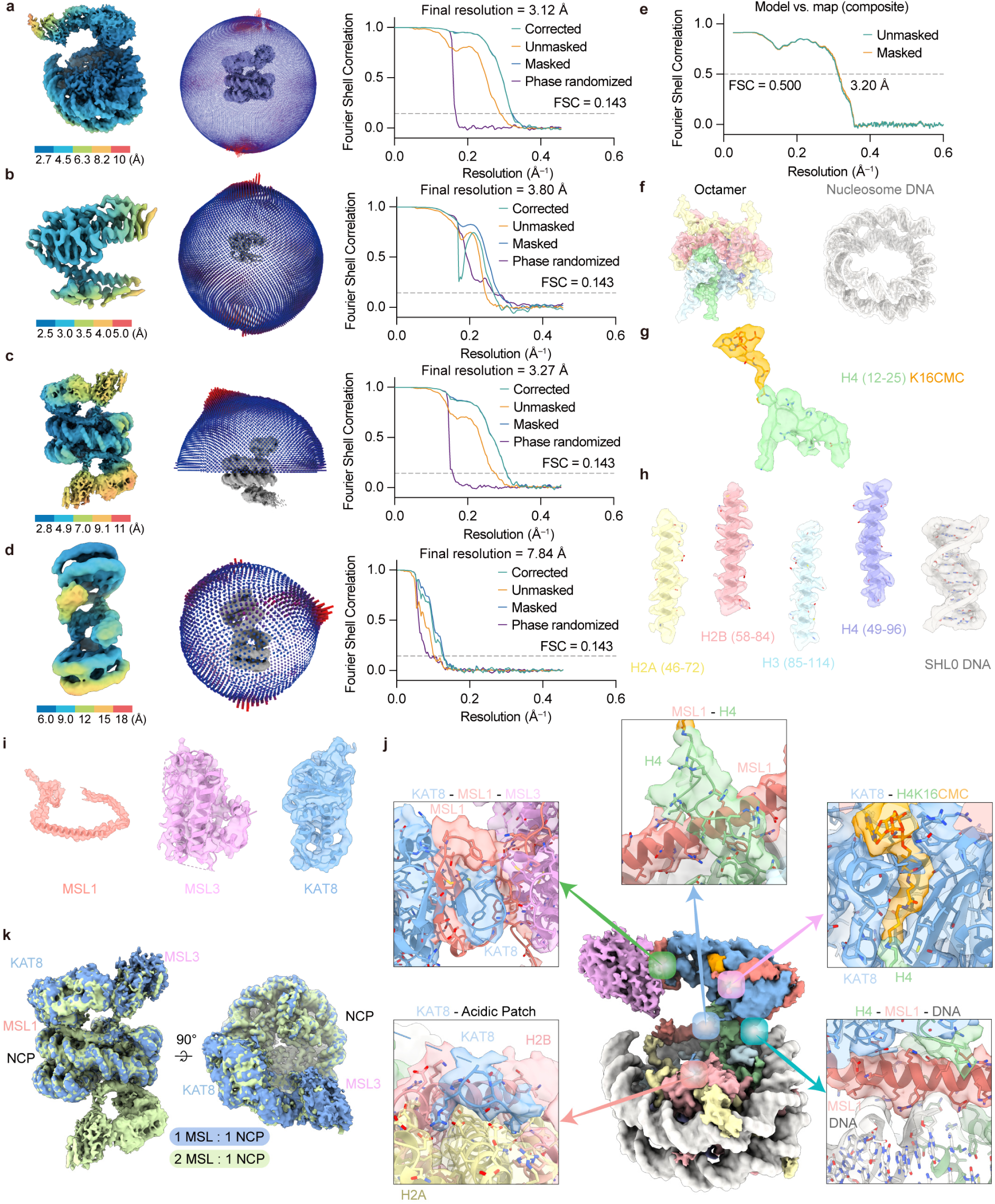
Cryo-EM map validation and interface analysis of the MSL–H4K16CMC nucleosome complex. **a**–**d**, The final reconstruction colored by local resolution, Euler angle distributions of the final reconstruction and Fourier shell correlation curve calculated from final refinement for (**a**) MSL–NCP complexes at 1:1 stoichiometry, (**b**) local MSL1–KAT8 components, (**c**) MSL–NCP complexes at 2:1 stoichiometry and (**d**) MSL–NCP complexes at 2:2 stoichiometry. The resolution at FSC = 0.143 was highlighted. (**e**) Fourier shell correlation curve for model-map correlation shown at FSC = 0.5. **f**–**i**, Cryo-EM density superimposed with individual components of the MSL–H4K16CMC nucleosome complex. (**f**) Histone octamer and nucleosome DNA. (**g**) H4 (12–25) segment containing a CMC-conjugated K16 side chain. (**h**) Selected regions of core histones. (**i**) MSL1, MSL3, and KAT8 subunits of MSL complex. **j**, Cryo-EM density of key interfaces within the MSL–nucleosome complex: the MSL1–H4 interface, the KAT8–H4K16CMC interface, the H4–MSL1–DNA interface, the KAT8–H2A/H2B acidic patch interface, and the KAT8–MSL1–MSL3 interface. Cryo-EM density is shown superimposed with the corresponding atomic models. **k**, Structural comparison of MSL–nucleosome assemblies at 1:1 and 2:1 stoichiometries.

**Extended Data Fig. 4.**
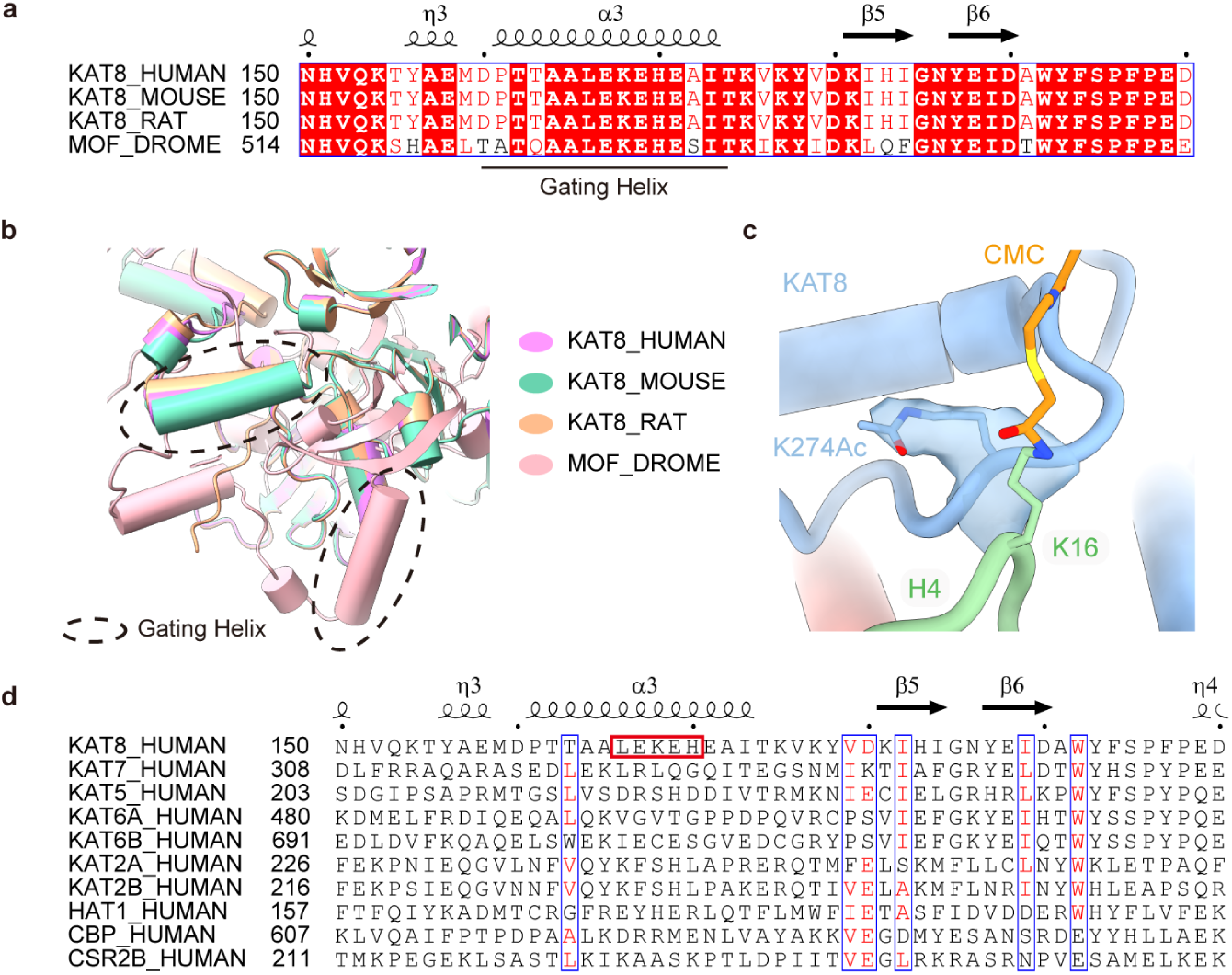
Conserved and specific KAT8 gating elements underlie H4-tail engagement in the MSL–nucleosome complex. **a**, Multiple sequence alignment of the gating-helix region of KAT8 orthologs. Secondary-structure elements are indicated above the alignment, with the gating helix underlined **b**, Structural superposition of KAT8/MOF orthologs from AlphaFold3 predictions highlighting the position of the gating helix (dashed outlines). In contrast to the distinct positioning observed in Drosophila MOF, vertebrate KAT8 orthologs adopt a conserved gating-helix position. **c**, Cryo-EM density map showing Ac-K274 of KAT8 gating hairpin, with the side chain fitted into the corresponding density. **d**, Multiple sequence alignment of the gating-helix region of human KAT8 with representative human histone acetyltransferases. Secondary-structure elements are indicated above the alignment; residues within the gating helix that contact histone H4 are highlighted by red boxes.

**Extended Data Fig. 5.**
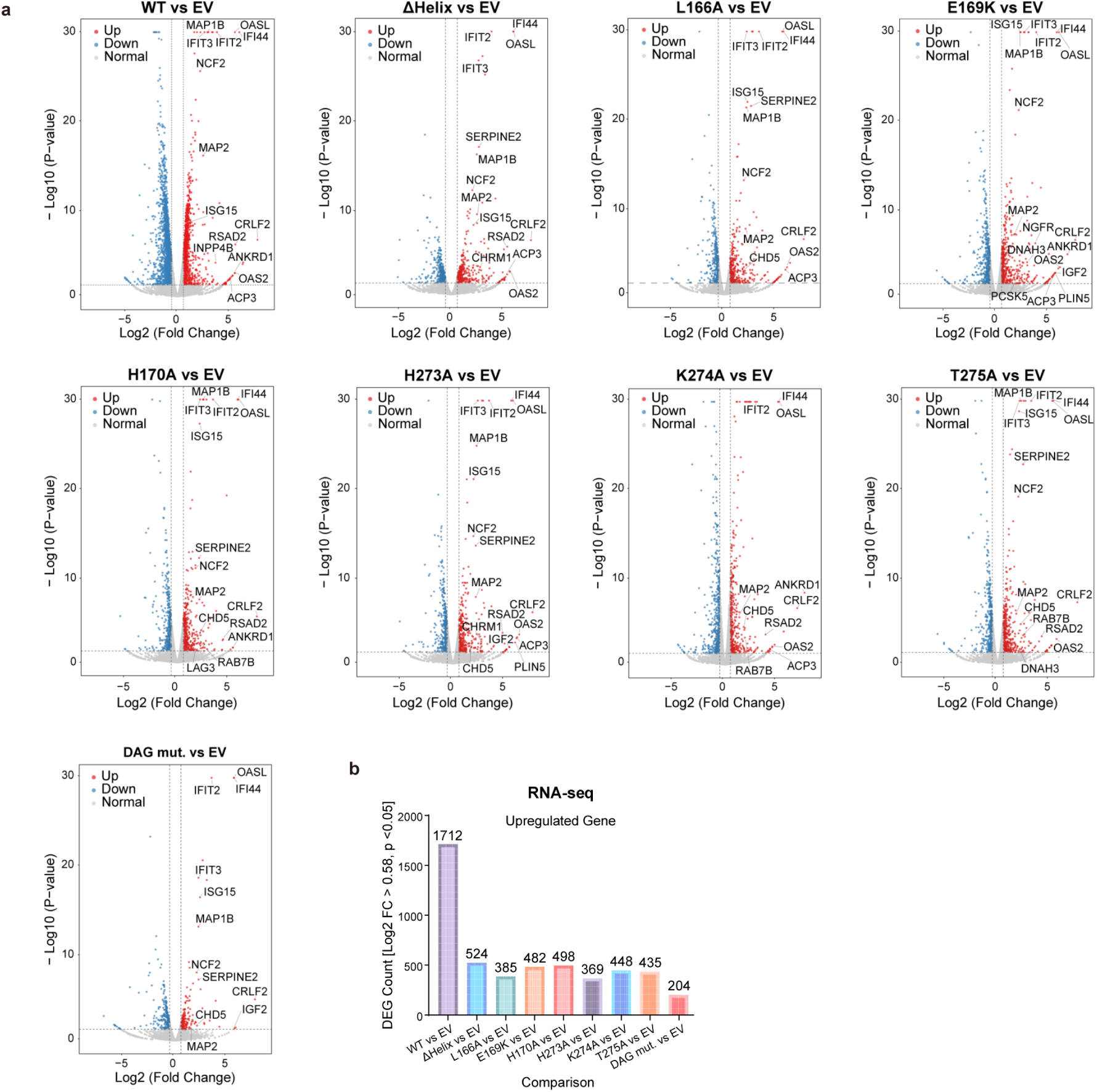
RNA-seq analysis reveals distinct gene expression defects caused by KAT8 mutations. **a**, Volcano plot of differentially expressed genes in *KAT8* KD cells expressing the indicated KAT8 constructs. **b**, Number of downregulated genes differentially expressed in *KAT8* KD cells expressing the indicated KAT8 constructs.

**Extended Data Fig. 6.**
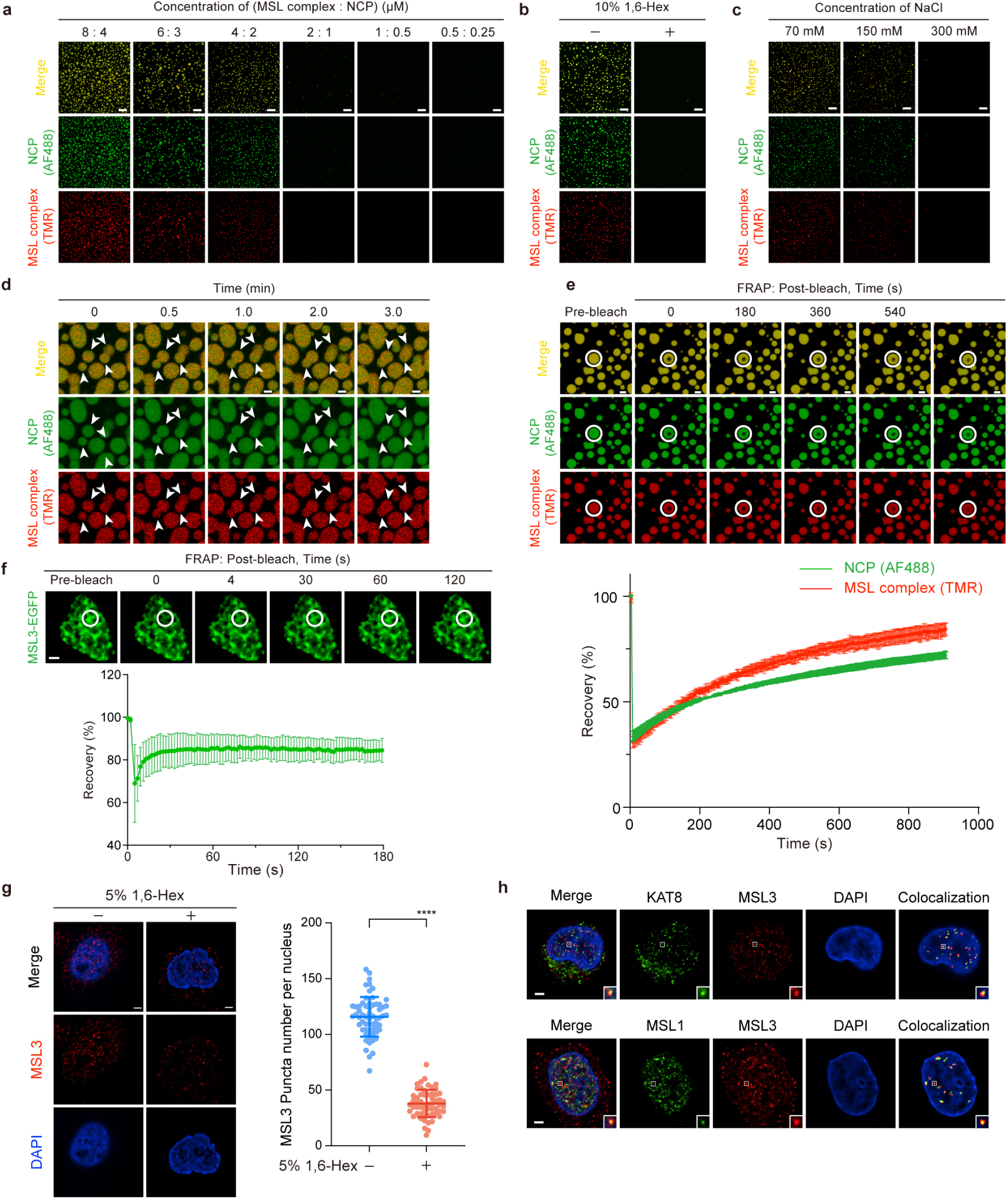
Dynamic phase separation mediated by multivalent MSL3 interactions. **a**, Representative fluorescence images of condensates formed by mixing MSL complexes with NCP in vitro at the indicated concentrations. Scale bar, 20 μm. **b**, Representative fluorescence images of MSL–NCP condensates in vitro treated with or without 10% 1,6-Hex. Scale bar, 20 μm. **c**, Salt dependence of MSL–NCP condensate in vitro formation at the indicated NaCl concentrations. Scale bar, 20 μm. **d**, Arrows indicate representative condensates formed by MSL complex and NCP in vitro, illustrating droplet fusion over the indicated time course. Scale bar, 2.5 μm. **e**, FRAP analysis of MSL–NCP condensates in vitro; Data are represented as mean ± s.d. from n = 4 biological independent experiments. Scale bar, 2.5 μm. **f**, FRAP analysis of MSL3–EGFP condensates in cells; Data are represented as mean ± s.d. from n = 11 biological independent experiments. Scale bar, 2 μm. **g**, Left, representative immunofluorescence images of nuclear puncta formed by endogenous MSL3 in cells treated with or without 5% 1,6-Hex. Right, quantification of nuclear MSL3 puncta per nucleus; n = 61 cells without 5% 1,6-Hex treatment and n = 57 cells with 5% 1,6-Hex treatment. A two-sample, two-tailed Student’s t-test was employed to calculate P values; ****p < 0.0001. Data are represented as mean ± s.d. Scale bar, 2 μm. **h**, Representative immunofluorescence images showing colocalization of MSL3 puncta with KAT8 or MSL1 in the nucleus. Scale bar, 2 μm.

**Extended Data Fig. 7.**
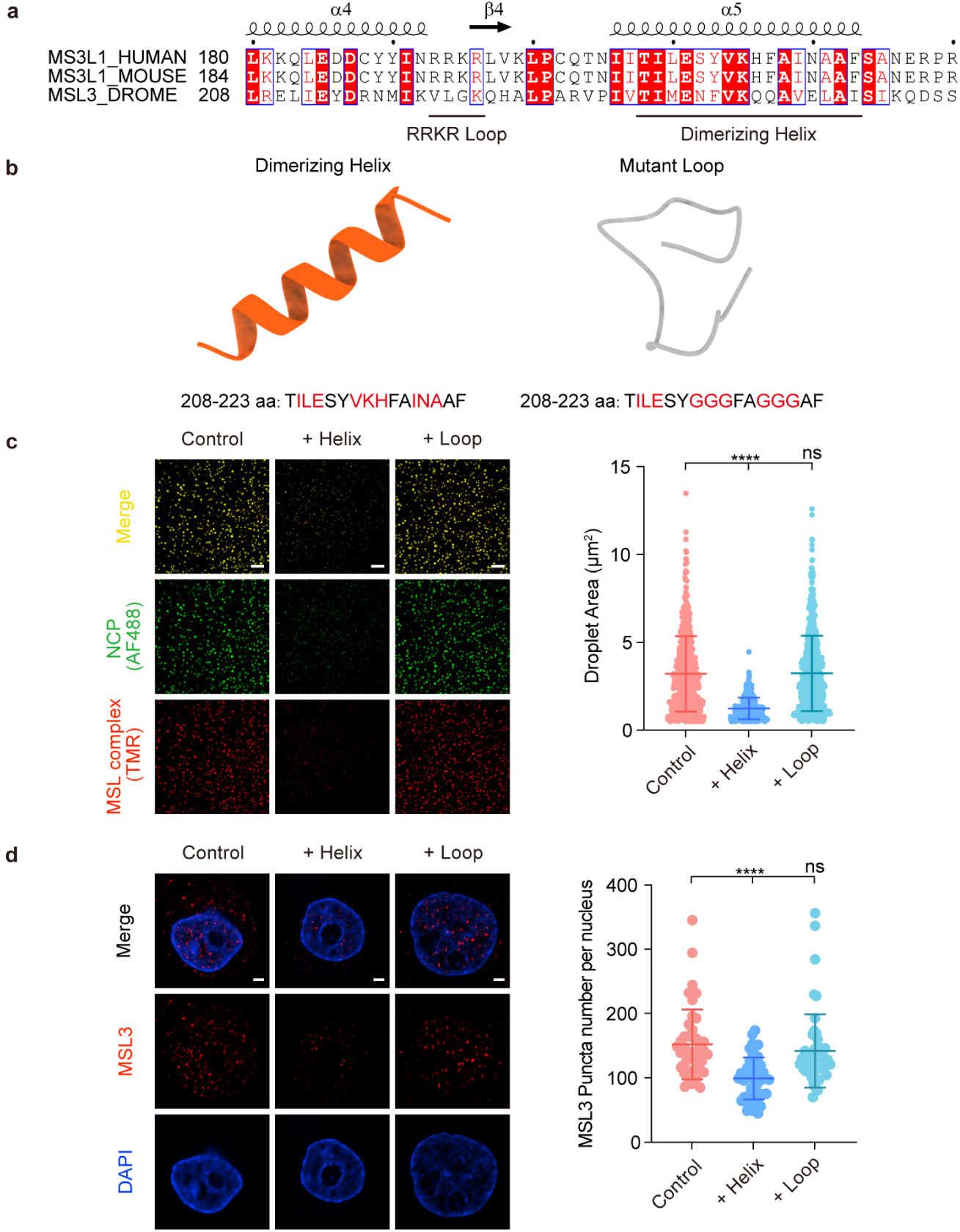
Chemical perturbation of the MSL3 dimerization interface disrupts phase separation. **a**, Multiple sequence alignment of the RRKR loop and dimerizing helix regions of MSL3 orthologs. **b**, Structure and sequence of dimerizing helix and mutant loop peptides derived from MSL3 dimerization interface. **c**, Left, representative fluorescence images of in vitro condensates formed by MSL complexes mixed with NCPs in the presence of the MSL3 dimerizing helix peptide or mutant loop peptide. Scale bar, 20 μm. Right, quantification of the droplet area. **d**, Left, representative immunofluorescence images of nuclear puncta formed by endogenous MSL3 in cells treated with the MSL3 dimerizing helix peptide or mutant loop peptide. Scale bar, 2 μm. Right, quantification of nuclear puncta formed by endogenous MSL3 in cells treated with the MSL3 dimerizing helix peptide or mutant loop peptide. n = 44 cells without peptide treatment, n = 47 cells with MSL3 dimerizing helix peptide treatment, and n = 50 cells with mutant loop peptide treatment. In (**c**) and (**d**), A two-sample, two-tailed Student’s t-test was employed to calculate P values; ****p < 0.0001 and ns means not significant. Data are represented as mean ± s.d.

**Extended Data Fig. 8.**
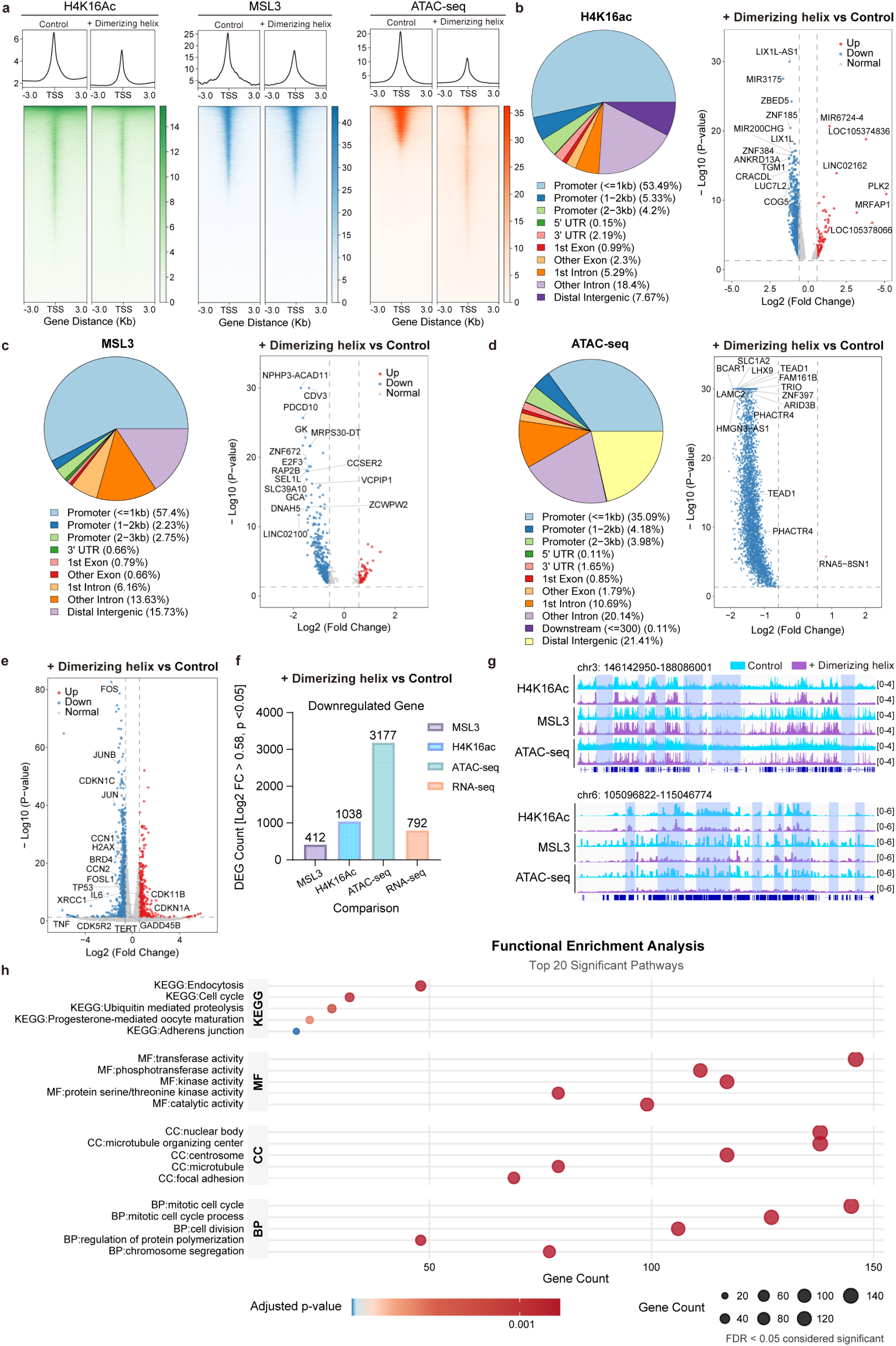
Chemical perturbation of MSL3-mediated phase separation alters downstream chromatin regulation and cellular functions. **a**, Heatmaps and average signal profiles of H4K16ac and MSL3 CUT&Tag and ATAC-seq signal in control cells and cells treated with MSL3 dimerizing helix peptide. **b**–**d**, Genome-wide distribution of differential H4K16ac-binding peaks, MSL3-binding peaks and chromatin accessibility regions and volcano plots of differential genes in control cells and cells treated with MSL3 dimerizing helix peptide. **e**, Volcano plots of differential genes in RNA-seq in control cells and cells treated with MSL3 dimerizing helix peptide. **f**, Numbers of downregulated genes differentially expressed between control cells and cells treated with MSL3 dimerizing helix peptide in multiple omics analyses. **g**, Genome browser views of representative loci illustrating coordinated spreading changes in H4K16ac, MSL3 occupancy and ATAC-seq signal in control cells and cells treated with MSL3 dimerizing helix peptide. **h**, Functional enrichment analysis of the reduced H4K16ac binding genes, significantly enriched KEGG pathways and Gene Ontology terms for control cells and cells treated with MSL3 dimerizing helix peptide.

**Extended Data Fig. 9.**
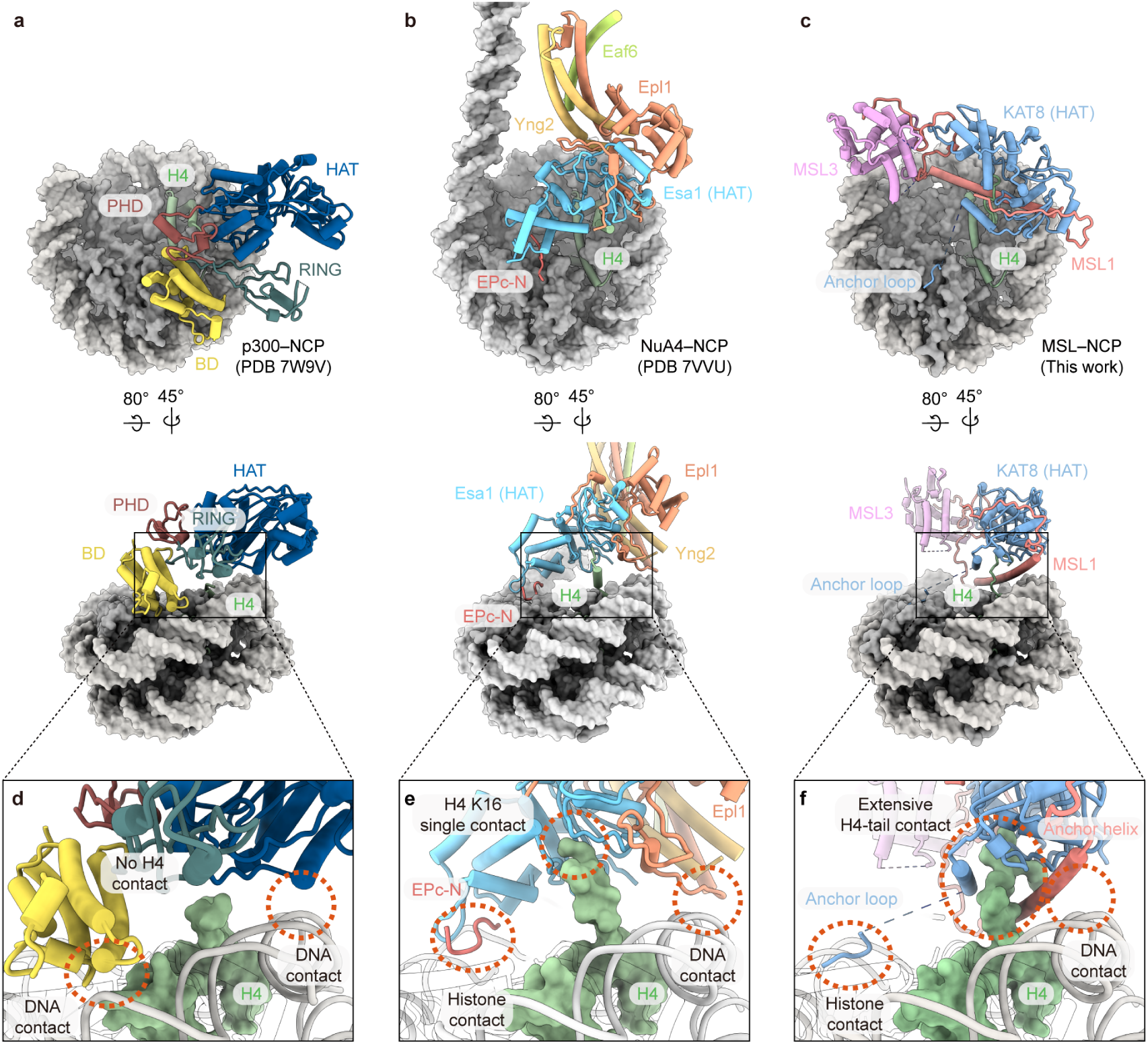
Distinct structural mechanisms govern histone acetylation modes on the nucleosome. **a**, Structure of p300 bound to the nucleosome (PDB 7W9V) shown in two views. **b**, Structure of the NuA4 HAT acetyltransferase module bound to the nucleosome (PDB 7VVU) shown in two views. **c**, Cryo-EM structure of the human MSL–nucleosome complex (this work) shown in two views. **d**–**f**, Close-up views of the boxed regions in (**a**) to (**c**), illustrating distinct organizational principles by which acetyltransferases engage the nucleosome: **d**, a DNA-centered binding mode lacking histone anchoring by p300; **e**, a position-based association that places the catalytic core near H4K16 without enforcing a fixed H4 trajectory by NuA4; **f**, a hierarchical integration of catalytic and non-catalytic contacts that constrain a single productive H4-tail trajectory by MSL complex.

**Extended Data Fig. 10.**
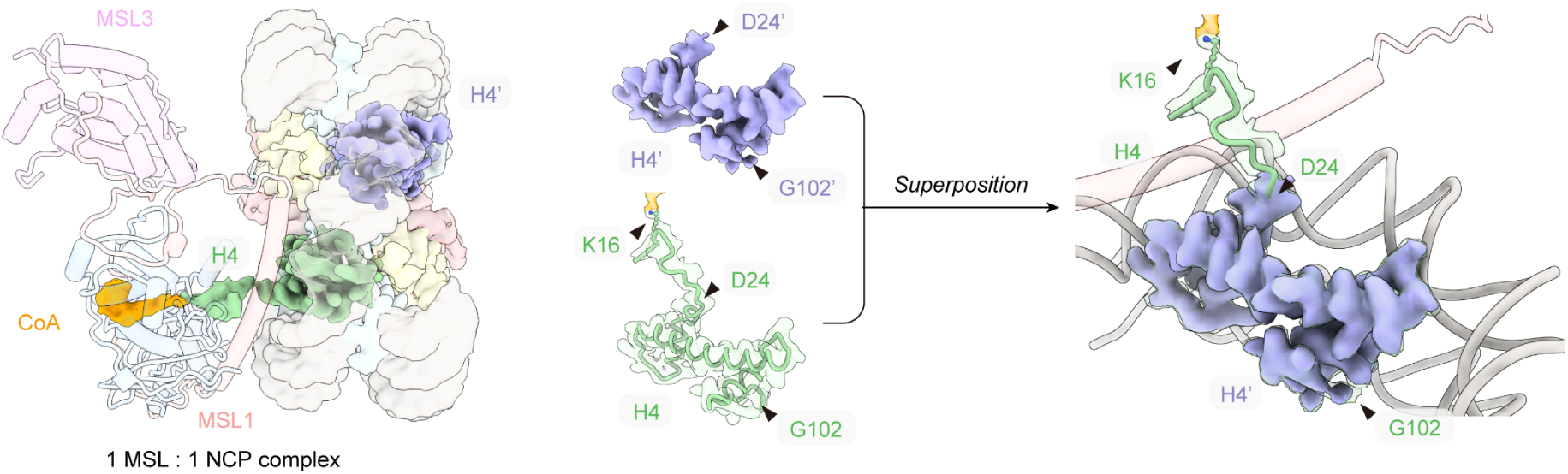
Chemical probe–assisted stabilization of a catalytically engaged MSL–NCP conformation. Comparison of H4 N-terminal tail conformations on the MSL-engaged and opposite nucleosomal faces within the 1:1 MSL–nucleosome complex. H4 N-tail density is shown in green for the MSL-engaged face and in purple for the opposite face. Superposition reveals enhanced stabilization and ordered positioning of the H4 tail under catalytic-engaged conditions enabled by the chemical probe, with increased visibility of the N-terminal H4 tail region preceding D24.

