## Supplementary File for "Hierarchical Anchoring–Gating Dictates Specific H4K16 Acetylation by the Human MSL Complex"

#### Supplementary Data 1

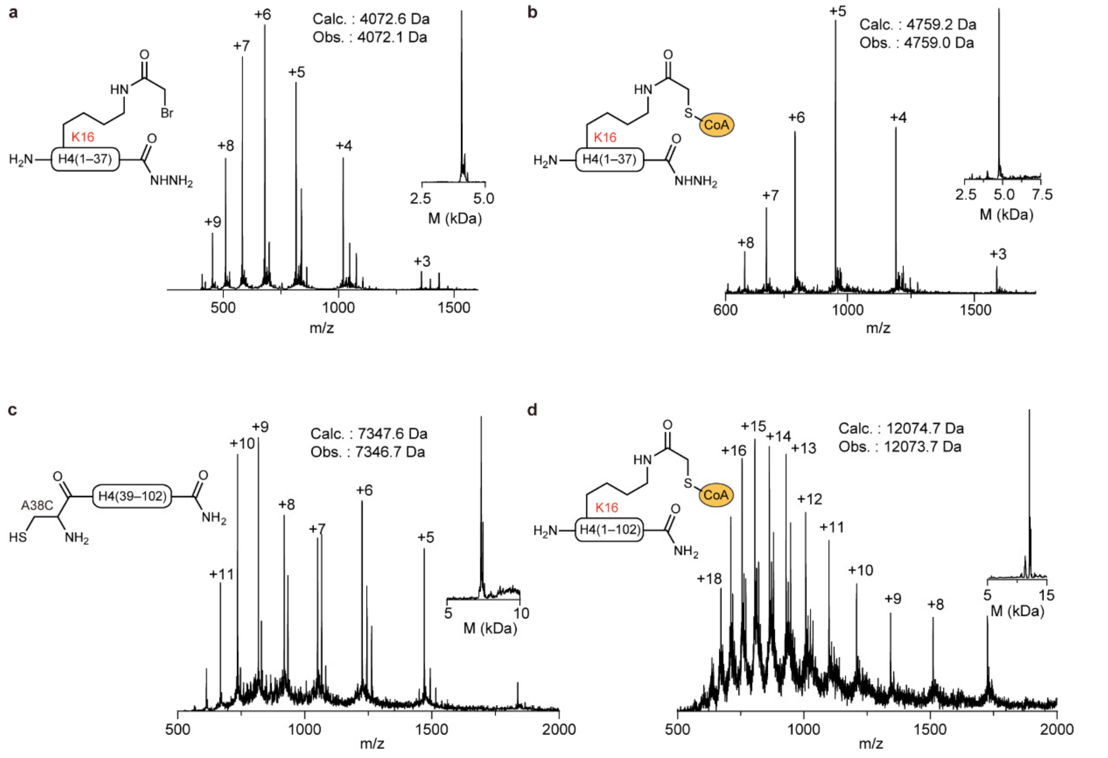


**Supplementary Data 1. Mass spectrometry of H4K16CMC probe synthetic intermediates. a–d**, Representative ESI-MS spectra and deconvoluted MS spectra of synthetic intermediates**.**

#### Supplementary Data 2: Uncropped blot images in the article

For Main Figures

**Fig. 1b**

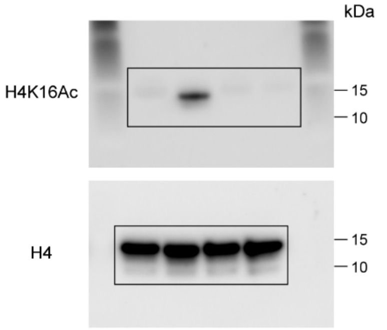


**Fig. 1c**

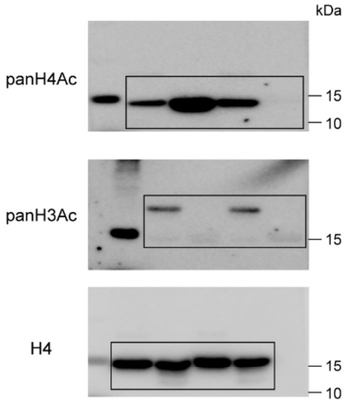


**Fig. 1d**

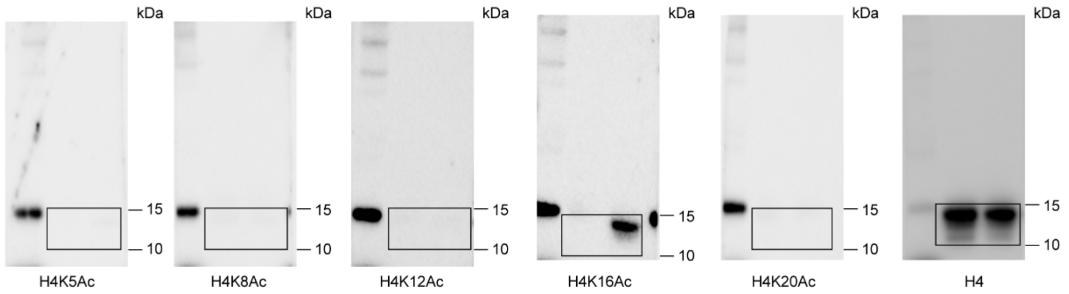


**Fig. 2f**

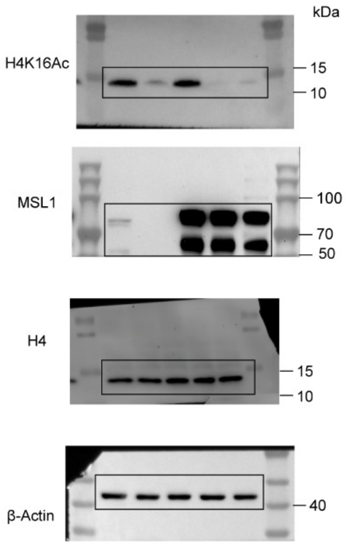


**Fig. 2g**

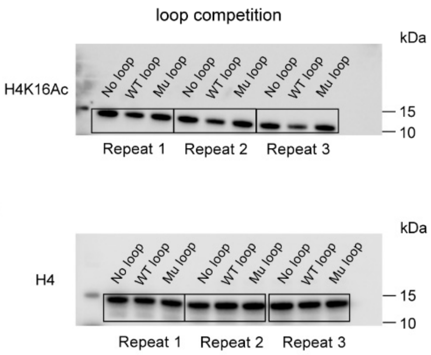


**Fig. 2i**

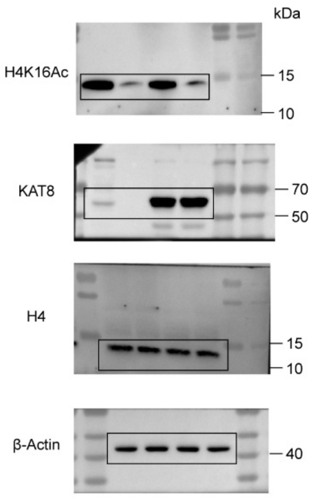

### Supplementary Data 2: Uncropped blot images in the article

#### For Main Figures

**Fig. 3i**

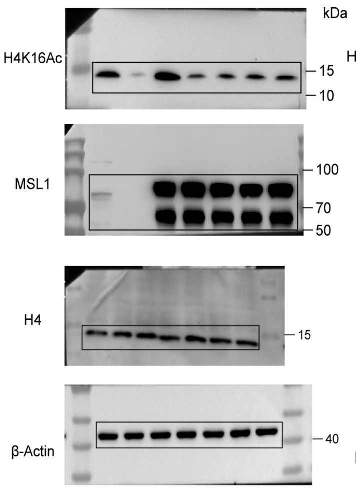


**Fig. 3j**

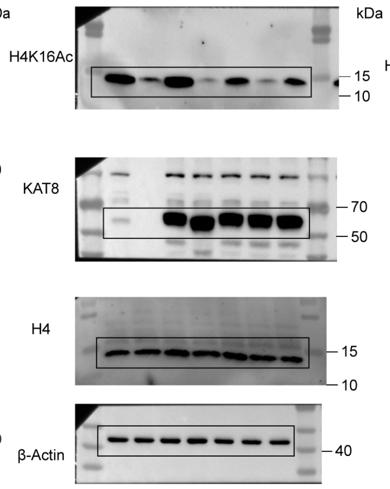


**Fig. 3k**

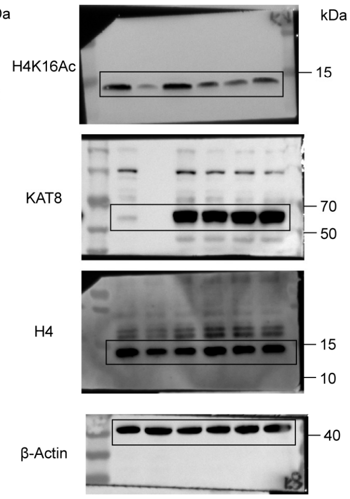


#### For Extended Data Figures

**Extended Data Fig.1b**

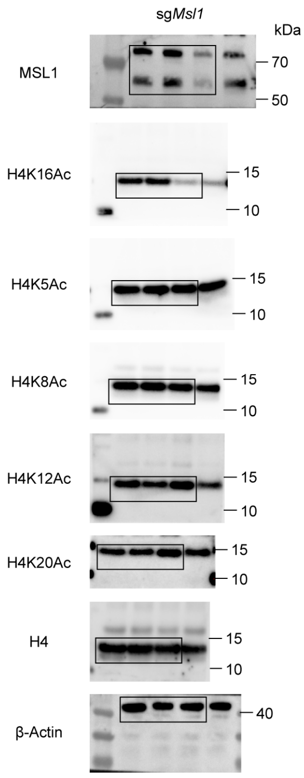


**Extended Data Fig.1c**

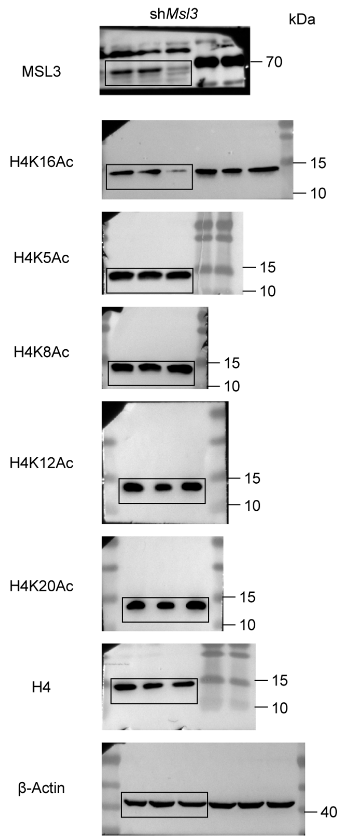


**Extended Data Fig.1d**

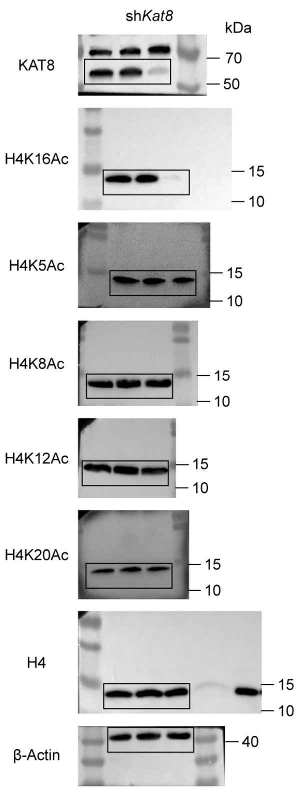

**Supplementary Table 1. Cryo-EM data collection, refinement, and validation statistics for MSL – NCP structures.**

| | MSL–NCP structures | | |
|---|---|---|---|
| **Stoichiometry** | **1:1** | **2:1** | **2:2** |
| **PDB entry** | 23FT | | |
| **EMDB entry** | EMD-68933 | EMD-68934 | EMD-68936 |
| **Data collection & processing** | | | |
| **Microscope and detector** | | K3 | |
| **Magnification** | | 64,000 | |
| **Voltage (keV)** | | 300 | |
| **Electron exposure ($e^-$/Å$^2$)** | | 50, 32 frames | |
| **Defocus range (μm)** | | –1.0 to –2.0 | |
| **Pixel size (Å)** | | 1.0979 | |
| **Micrographs (no.)** | | 9,923 | |
| **Symmetry imposed** | C1 | C2 | C1 |
| **Final particles (no.)** | 245,245 | 174,261 | 12,631 |
| **Map global resolution (Å)** | 3.12 | 3.27 | 7.84 |
| **FSC threshold** | 0.143 | 0.143 | 0.143 |
| **B-factor (Å$^2$)** | –30 | –30 | –100 |
| **Refinement & validation** | | | |
| **Initial model** | PDB 7VVU, 7XD1, 2Y0N, 2Y0M; Alphafold3 prediction | | |
| **Model resolution (Å)** | 3.20 | | |
| **FSC threshold** | 0.5 | | |
| **Model composition** | | | |
| **Non-hydrogen atoms** | 16,988 | | |
| **Protein residues** | 1,350 | | |
| **Nucleotides** | 294 | | |
| **R.M.S. deviations** | | | |
| **Bond lengths (Å)** | 0.004 | | |
| **Bond angles (°)** | 0.594 | | |
| **B-factor (Å$^2$)** | | | |
| **Protein (min/max/mean)** | 30.00/1319.95/287.47 | | |
| **Nucleotides (min/max/mean)** | 69.18/221.90/128.08 | | |
| **MolProbity score** | 1.44 | | |
| **Clashscore** | 6.87 | | |
| **Poor rotamers (%)** | 0.00 | | |
| **CaBLAM outliers (%)** | 0.93 | | |
| **Ramachandran plot** | | | |
| **Favored (%)** | 97.73 | | |
| **Allowed (%)** | 2.27 | | |
| **Outliers (%)** | 0.00 | | |

**Supplementary Table 2. shRNA and sgRNA oligonucleotides used in this study.**

| Target | Sequence |
|---|---|
| **KAT8** | shRNA-1: TGTACGAAGTTGATGGCAAAG |
| | shRNA-2: CAGAAGAACTCAGAGAAGTAC |
| **MSL3** | shRNA-1: GGCAGATCATCTTCACCTAT |
| | shRNA-2: GCCAACATGAACGTGCATTA |
| **MSL1** | sgRNA: GGATTGAACGTATGGAAAGG |
